# Ancient whole genome duplications and the evolution of the gene duplication and loss rate

**DOI:** 10.1101/556076

**Authors:** Arthur Zwaenepoel, Yves Van de Peer

## Abstract

Gene tree - species tree reconciliation methods have been employed for studying ancient whole genome duplication (WGD) events across the eukaryotic tree of life. Most approaches have relied on using maximum likelihood trees and the maximum parsimony reconciliation thereof to count duplication events on specific branches of interest in a reference species tree. Such approaches do not account for uncertainty in the gene tree and reconciliation, or do so only heuristically. The effects of these simplifications on the inference of ancient WGDs are unclear. In particular the effects of variation in gene duplication and loss rates across the species tree have not been considered. Here, we developed a full probabilistic approach for phylogenomic reconciliation based WGD inference, accounting for both gene tree and reconciliation uncertainty using a method based on the principle of amalgamated likelihood estimation. The model and methods are implemented in a maximum likelihood and Bayesian setting and account for variation of duplication and loss rate across the species tree, using methods inspired by phylogenetic divergence time estimation. We applied our newly developed framework to ancient WGDs in land plants and investigate the effects of duplication and loss rate variation on reconciliation and gene count based assessment of these earlier proposed WGDs.

## Introduction

In the past decades, examination of genomic data has revealed many signatures of ancient whole genome duplications (WGDs) across the eukaryotic tree of life (reviewed in Van de Peer et al. 2017). These findings have initiated an active research field concerned with the evolutionary importance of polyploidy, especially in plants where polyploidization seems to have been rampant. The apparent widespread incidence of polyploidy in the phylogeny of land plants is often cited as strong evidence for the evolutionary importance of polyploidy. But just how widespread is ancient polyploidy in land plants? While there has been strong evidence for many (relatively recent) WGD events, inference of these events, especially very ancient ones, remains highly challenging. Evidently, the signal of an ancient WGD event erodes through time, and all current methods suffer a strong loss of power the more ancient the hypothesized event. As a result, some of the claimed ancient WGD events have been contested, such as the two hypothesized events in land plants (Jiao et al. 2011; Ruprecht et al. 2017), the 2R hypothesis in vertebrates (Abbasi 2010; Van de Peer et al. 2010; Smith and Keinath 2015), or ancient WGD events in hexapods (Zheng Li et al. 2018; Li et al. 2019; Nakatani and McLysaght 2019).

Methods for unveiling ancient WGDs can be classified crudely into three main approaches. The first approach takes advantage of the expectation that a WGD leaves a signature in the distribution of duplicate divergence times. One commonly estimates the synonymous distance (*K_S_* or d*S*, which serves as a proxy for the divergence time) for all paralogous pairs in a genome and visualizes the resulting distribution. In such a *K_S_* distribution, ancient WGDs will be visible as peaks against the background exponential decay distribution from small scale duplication (SSD) events (Lynch and Conery 2000; Blanc and Wolfe 2004). There are a couple of pitfalls with this approach, which have been discussed in detail (Vanneste et al. 2013; Tiley et al. 2018; Zwaenepoel et al. 2018). Importantly, these distributions are not suitable for inferring very ancient events due to saturation of the synonymous distance. The second main approach is based on the expectation that a WGD should lead to large co-linear blocks in the genome. Such co-linearity or synteny based information has often been considered as the strongest evidence for ancient WGDs. In particular the combination of syntenic and *K_S_* information has been vital for the discrimination of WGD-derived and SSD-derived paralogs. The major drawback is however that high quality genome assemblies are required, and these are still non-trivial to obtain. Nevertheless, even with high quality assemblies, interpretation of syntenic signal for very ancient putative WGDs is not always unequivocal. In particular the temporal (either relative or absolute) framing of a WGD event based on syntenic data is complicated and requires high quality genomes of multiple related lineages. The last set of methods are united by their usage of phylogenetic information in individual gene families. Both methods using gene counts and gene tree topologies have been used, either in a model-based or heuristic framework. Especially heuristic gene tree - species tree reconciliation methods have been widely employed to unveil evidence for ancient WGDs (e.g. Jiao et al. 2011; Li et al. 2015; Li et al. 2016; McKain et al. 2016; Thomas et al. 2017; Zheng Li et al. 2018; Yang et al. 2018), often in combination with other sources of evidence. Here, we take heuristic to mean that the gene tree is inferred independently from its reconciliation. In these approaches, a larger than expected number of duplication events inferred for a particular branch of the species tree is regarded as indicative for an ancient WGD. So far, most of the support for very ancient WGD events has been obtained using gene tree reconciliation approaches. These approaches naturally provide a temporal view on the hypothesized event as they assume a known, either dated or undated, species tree.

There are however several potential pitfalls when employing heuristic gene tree - species tree reconciliation approaches. The first, and probably most obvious, is the need for an arbitrary cut-off on the number of duplications before some species tree branch is associated with a WGD. This is especially troubling for putative WGD events on tip branches, as large numbers of SSD events can easily be confused with a WGD event (Zwaenepoel et al. 2018). The number of duplication events inferred for specific branches can also be very sensitive to taxon sampling, and some signal for a putative WGD event on a particular branch may be absent or weakened when the branch is subdivided by adding more taxa to the analysis. Perhaps more important are the problems with the methodology *per se*. In most cases, reconciliation approaches rely on a single gene tree topology for every gene family, inferred by maximum likelihood methods, and a single reconciliation thereof, typically employing a least common ancestor (LCA) approach which minimizes the total number of duplication and loss events (Zmasek and Eddy 2001). A gene tree topology is however a probabilistic model of the phylogeny of that gene family, and for a single gene family there may be a considerable number of different topologies with near equal support (Salter 2001). Similarly, a reconciliation of a gene tree to a species tree can also be considered probabilistically, and, although less well studied, relying on the single most parsimonious reconciliation may be similarly problematic. In particular the joint effects of these two issues may be of crucial importance, as the reconciliation of uncertain topologies by means of LCA reconciliation will result in conflicting views on the evolution of the gene family and systematic biases (see e.g. Hahn 2007). To overcome some of these problems, researchers have typically filtered out nodes with low bootstrap support, evaluated some type of duplication consistency scores or have used heuristic branch swapping methods in the reconciliation step (as implemented for example in Notung (Chen et al. 2000)).

Probabilistic methods for WGD inference in a phylogenetic context, both employing gene trees and gene counts, were recently proposed by Rabier et al. (2014). In a gene count based method (Hahn et al. 2005), one does not employ topological information but effectively integrates over all possible gene trees that could have generated the observed counts at the species tree leaves. Such methods therefore naturally handle uncertainty in the gene tree, albeit in a somewhat crude fashion. The observed gene family is modeled as the outcome of a birth-death Markov chain, allowing likelihood based inference of duplication and loss rates, ancestral gene counts and, in the framework of Rabier et al. (2014), WGD retention rates. While they have yielded great insights in genome evolution, gene count based methods do not consider all of the information in genomic data sets, as sequence data for the genes provides information about their phylogeny. Therefore, a gene tree - species tree reconciliation approach that estimates parameters of a model of gene family evolution is expected to be more accurate. We expect this in particular when models are employed that allow variation in the duplication and loss rate across the species tree. Additionally, reconciliation based methods have the obvious advantage of providing the researcher with an actual reconciled gene tree, i.e. a tree with nodes labeled as either a speciation, duplication or loss node. In our case, this labeling should also include whether a particular duplication node is inferred to be a WGD or SSD-derived duplication. This therefore also provides a model-based framework for selecting gene families for Bayesian molecular dating analyses to estimate absolute ages of ancient WGDs (as in e.g. Vanneste et al. 2014; and Clark and Donoghue 2017) or to study functional biases in gene retention patterns (e.g. Li et al. 2016). Contrary to expectations, Rabier et al. (2014) reported a lower power to detect WGDs for their reconciliation approach compared to their gene count approach, and they recommend usage of the gene count method for testing WGD hypotheses in a phylogenetic context. However, they attributed these observations mainly to computational limitations in their reconciliation method.

Here we introduce a novel method for WGD inference using gene trees designed to overcome the issues of the reconciliation method in Rabier et al. (2014). We draw insipiration from the growing body of literature on gene tree inference under a known species tree (reviewed in Szöllősi et al. 2015). Our approach is based on the principle of amalgamated likelihood estimation (ALE) for probabilistic gene tree - species tree reconciliation, first proposed and developed by Szöllősi, Rosikiewicz, et al. (2013). We develop an ALE approach, called Whale, employing the probabilistic model of Rabier et al. (2014) to estimate duplication, loss and WGD retention rates and test WGD hypotheses in a phylogenetic context. By using the amalgamation principle with a probabilistic model of gene family evolution in the presence of WGDs, Whale jointly accounts for uncertainty in the gene tree topology and reconciliation. As in Szollosi, Rosikiewicz, et al. (2013), our approach is fully probabilistic, and does not employ parsimony-guided reconciliation as in Rabier et al. (2014). We employed the Whale method both in a maximum likelihood and Bayesian setting and reveal the crucial importance of considering duplication and loss rate heterogeneity across the species tree when assessing WGD hypotheses. To accommodate this, we implemented models of duplication and loss rate evolution inspired by molecular divergence time estimation. Revisiting some of the ancient WGDs reported in the land plant phylogeny, we evaluated our new approaches and discuss caveats when assessing WGDs using gene tree reconciliation.

## New approaches

- We implemented algorithms to compute the joint gene tree - reconciliation likelihood under the probabilistic model of Rabier et al. (2014) using the principle of amalgamation.
- Through analysis of simulated and empirical data sets we show that likelihood based inference of whole genome duplications (WGDs) *sensu* Rabier et al. (2014) is very sensitive to rate variation across branches of the species tree. This also has implications for simulation-based assessment of putative ‘bursts’ in the number of duplications in data sets of reconciled gene trees.
- We implemented models that can accommodate variation in duplication and loss rates inspired by Bayesian divergence time estimation and employ these to study the evolution of the duplication and loss rate together with putative ancient WGDs.

## Results

### Validation using simulated data

The ALE approach and the dynamic programming algorithm for probabilistic reconciliation inference have been extensively validated using simulations (Szöllősi et al. 2012; Szöllősi, Rosikiewicz, et al. 2013; Szöllősi, Tannier, et al. 2013). However, our adoption of these methods is considerably different from these studies, which focused mainly on horizontal gene transfer and improved gene tree inference under a known species tree. We introduce the WGD model as well as the prior distribution on the number of lineages at the root first developed by Rabier et al. (2014) in the ALE context, and estimate duplication and loss rates not family-wise as in ALE (Szollosi, Rosikiewicz, et al. 2013), but across families similar to Rasmussen and Kellis (2011) and Rabier et al. (2014) (see methods). We verified the correctness of our new approach and its implementation using simulated data. Importantly, while the ALE approach takes gene tree uncertainty into consideration by employing samples from the posterior distribution for gene tree topologies, we simulated only a single unrooted gene tree topology per family, and do not consider gene tree uncertainty here. This was previously done already using extensive simulations in Szollosi, Rosikiewicz, et al. (2013), where the basic merits of an ALE approach were shown, and we do not revisit these highly computationally intensive simulation studies here. Note that all reported rate estimates are dependent on the time scale used in the species tree, which in our case is in units of 100 million years.

Numerical optimization of the likelihood under the basic constant-rates duplication-loss (DL) model (i.e. using a single duplication (*λ*) and loss (*μ*) rate for the full species tree) with a geometric prior distribution on the number of lineages at the root provides accurate maximum likelihood estimates (MLEs) for the simulated duplication and loss rates (Figure S1). In general, rates are estimated more accurately when the duplication and loss rate are similar whereas slight biases are observed when the rates are quite different. If the loss rate is higher than the duplication rate, both rates tend to be underestimated. If the duplication rate is higher than the loss rate, the duplication rate seems to be slightly overestimated. Not unexpected, our simulations suggest that the variance of the MLEs increases with the rate. Estimates of the duplication (*λ*) and loss (*μ*) rate are quite robust to the parametrization of the geometric prior distribution on the number of genes at the root (Figure S2). As expected, assuming a very low prior probability on multiple genes at the root (1/*η* ≈ 1) leads to overestimation of *λ* and underestimation of *μ*. Conversely, assigning a strong prior on multiple ancestral lineages (1/*η* ≫ 1) leads to an underestimation of *λ* and overestimation of *μ* to compensate for unobserved lineages assumed at the root. These observations hold for different simulated values of *η*. We note that when gene trees are inferred for orthogroups obtained with software such as OrthoFinder or OrthoMCL, it is reasonable to use a prior with *η* > 0.5, as by the very definition of orthogroup we expect there to be a single ancestral lineage at the root of the species tree.

Similar simulations were performed for the DL + WGD model. We simulated three WGD scenarios under a constant-rates DL model with various retention rates. The three simulated scenario’s were (1) an ancestral WGD in seed plants (long internal branch), (2) an ancestral WGD in gymnosperms (short internal branch) and (3) a WGD in the *Physcomitrella patens* lineage (tip branch). When the simulated loss rate was low, the retention rate is accurately estimated (Figure 1). When the loss rate was at the higher end, both the retention rate and the loss rate tended to be underestimated for the seed plant and gymnosperm WGD scenarios. Such biases in the estimated retention rate seem not to be present in the *P. patens* WGD scenario, however in this case the variance in the MLE for *q* is larger. No bias in the estimated duplication and loss rates conditional on the simulated retention rate was observed (Figure S3). Since loss from the linear birth-death process and non-retention after a WGD are confounded (i.e. one can not discriminate between both possibilities for individual loss events), a high loss rate together with a high retention rate provides a competing explanation to a low loss rate together with a low retention rate for the data on the WGD branch. The estimation of these rates is therefore expected to be sensitive to the topology and location of the hypothesized WGD, as well as the number of free parameters allowed (see below, where we consider variation in *λ* and *μ* across the species tree). This also relates to possible identifiability issues discussed further below.

We investigate the power and false positive rate (FPR) of the likelihood ratio test (LRT) proposed in Rabier et al. (2014) to test for the presence or absence of a WGD using 500 gene families. The LRT for the constant-rates analysis has very high power to detect a WGD for a data set of 500 gene families when simulated rates were constant across the species tree (Figure 2, *v* = 0.00). Whale seems to have considerable higher power than the gene count based method WGDgc developed by Rabier et al. (2014), and has a lower FPR (0% compared to 6%, based on 100 replicates) for this data set size. The accuracy of the estimated retention rates is comparable between the two methods, with a slightly larger spread for the estimates acquired with WGDgc. This shows, as was also expected by Rabier et al. (2014), that in the absence of model violations, a gene tree reconciliation based method has greater power than a gene count based method to detect WGDs. The Whale method therefore provides a possible solution to the problems with the reconciliation based method in Rabier et al. (2014).

**Figure 1:**
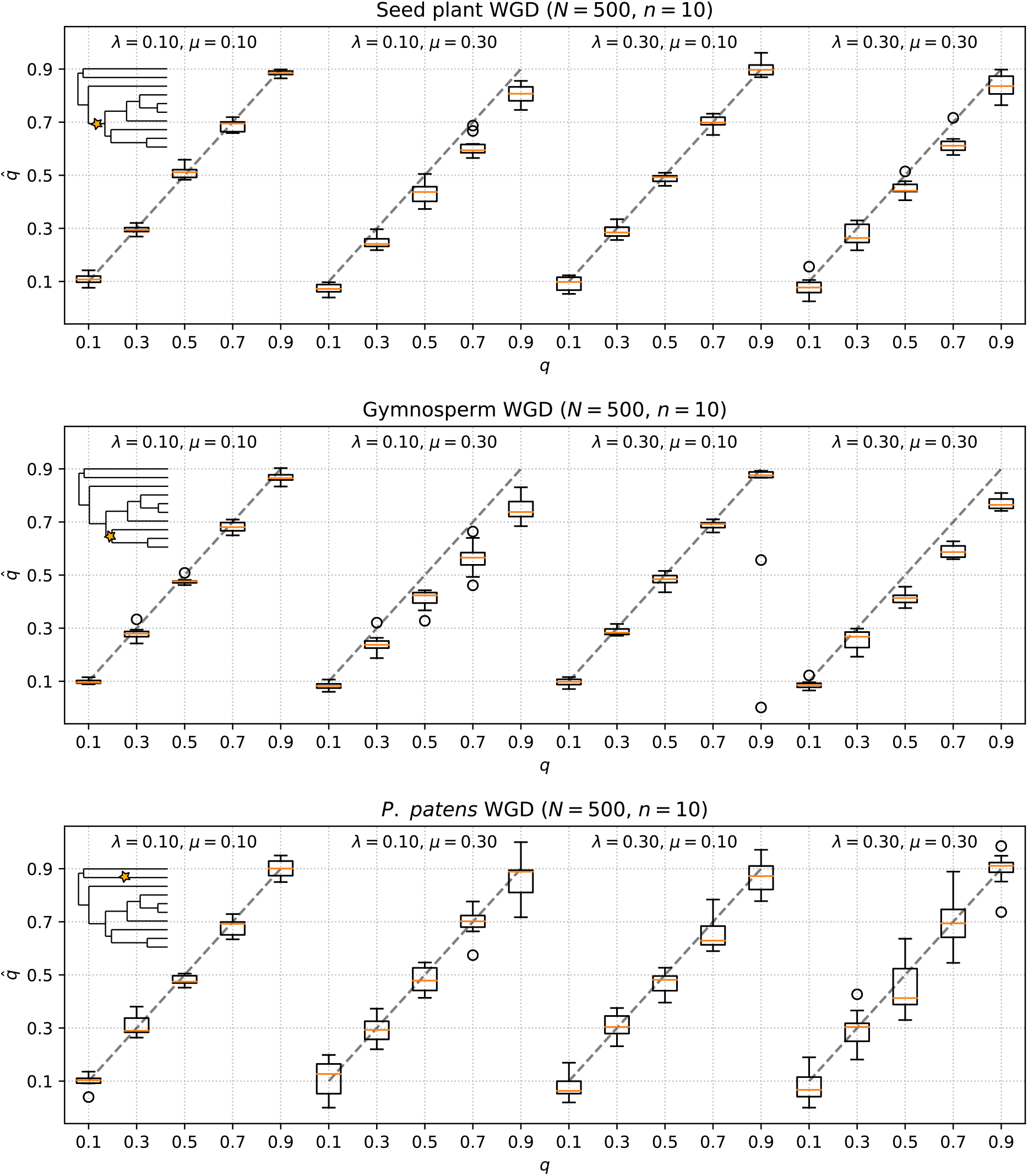
MLEs for the retention rate for different simulated WGD scenario’s and duplication and loss rates. Simulations of 10 times 500 gene families were done for a 10-taxon tree with constant duplication and loss rates across the tree. The species tree topology used is shown in the upper left corner of each set of simulations, and the star indicates the simulated WGD event.

**Figure 2:**
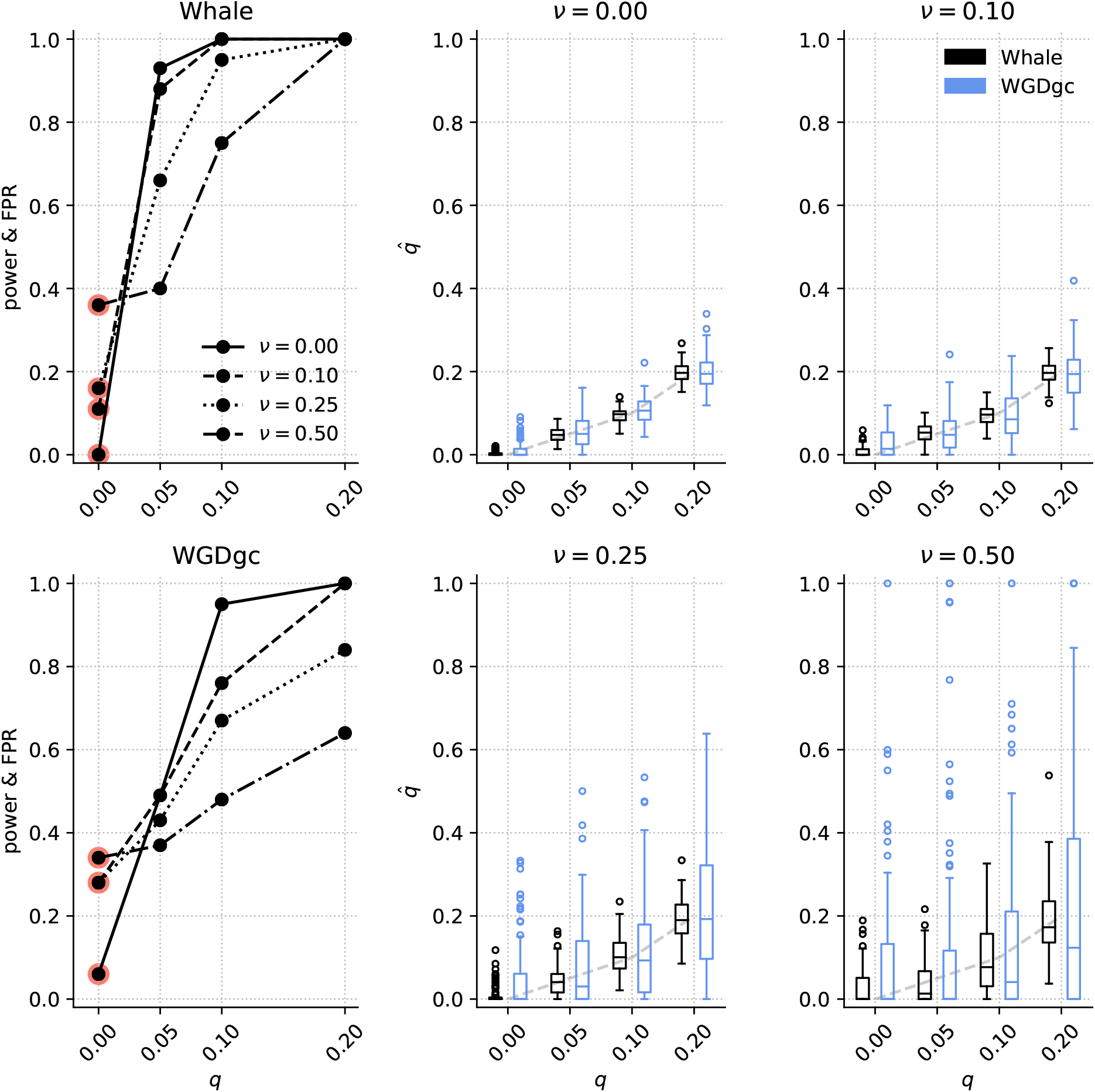
Power and false positive rate (FPR) of the LRT statistic to detect a simulated WGD and accuracy of the estimated retention rates for varying levels of rate heterogeneity. The two leftmost plots show the power and FPR (highlighted in red) of Whale and WGDgc based on 100 simulations of 500 gene families for every retention rate *q* and every level of rate heterogeneity *v*. The boxplots show the MLEs for the retention rates, with the MLEs from Whale in black and those from WGDgc in blue. The same 10-taxon tree was used as in Figure 1, with as simulated WGD the seed plant WGD event.

Similar to other molecular evolutionary rates, duplication and loss rates may vary across lineages of the species tree. The effect of such rate variation was not examined in Rabier et al. (2014), however we expect such rate variation to have an impact on the reliability and power of WGD inference. We simulated rate variation by drawing rates for every species tree branch from a log-normal distribution with mean 0.10 for *λ* and 0.15 for *μ* and standard deviation *v* for both. We put a lower and upper limit on both duplication and loss rates to retain realistic values (0.05 and 0.25 for *λ*, 0.10 and 0.50 for *μ*). The accuracy of the retention rate estimates drops with increasing rate variation (*v* > 0) and this is the case for both methods. Concomitantly, the power to detect a WGD drops quite dramatically, especially for WGDgc. More importantly, the FPR rises to inacceptable levels already for a relatively small amount rate variation (11% and 28% for Whale and WGDgc respectively with *v* = 0.10). Clearly, the LRT to test for WGD significance under the constant-rates DL + WGD model is not robust to model violations, and should not be used unless the hypothesis of variable rates across the species tree can be rejected. We note however that we cannot adopt a stage-wise approach where we first test the hypothesis of constant duplication and loss rates across the species tree before testing a WGD hypothesis, since the presence of *bona fide* WGD events will result in a rejection of the constant rates hypothesis. It is therefore crucial to account for rate variation across lineages and WGD events simultaneously, as our simulations show that failing to do so may seriously compromise the reliability of the LRT for testing a WGD hypothesis. We note that rate heterogeneity across gene families (as opposed to lineages) did not affect the FPR. Variability of duplication and loss rates across families has no considerable influence on the retention rate ML estimate (Figure S4).

### Maximum likelihood based inference for the land plant data set

We employed our approach for calculating the joint tree - reconciliation likelihood in a maximum likelihood (ML) framework to assess the evidence for several putative WGDs across the land plant phylogeny. We used a nine-taxon data set consisting of 7787 orthogroups, for each of which we obtained a sample from the posterior distribution of phylogenetic trees under the LG+Γ4 substitution model by means of MCMC (see methods). These samples were used to perform estimation of duplication, loss and retention rates with our newly developed framework. We focussed on evaluating hypotheses of very ancient WGDs, as relatively recent WGDs (up to about 100 Mya) can often be discerned from *K_S_* distributions or simpler reconciliation based analyses. In particular we focus on two widely accepted events on the branch leading to the seed plants (*ζ*) and the branch leading to the angiosperms (*ϵ*) (Jiao et al. 2011; Clark and Donoghue 2017). These hypothesized events have not been without controversy (Ruprecht et al. 2017), and it is therefore interesting to evaluate the evidence for these events under different models using our methods.

We first focus solely on the angiosperm and seed plant WGD. We adopt the approach proposed by Rabier *et al*. (2014), and tested extensively by Tiley *et al*. (2016), which consists of comparing the model fit of a DL+WGD model to a DL model by means of a likelihood ratio test (LRT). We set the prior on the root such that on average 1.5 genes are expected to be present at the root (*η* = 0.66). Using the LRT, we were unable to reject the null hypothesis of no WGD leading to the angiosperms (ML estimate for the retention rate 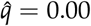), whereas a significantly better fit was obtained for a model with a WGD on the branch leading to the seed plants at 390 Ma (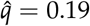, Λ = 1563.52, *p* < 0.001). The gene count based method of Rabier *et al*. (WGDgc) rejected both WGD hypotheses, with the MLE for both retention rates equal to zero, a result also obtained by Tiley et al. (2016). The MLEs for the birth and death rates were remarkably different, with 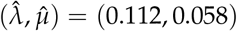 for WGDgc and (0.140,0.159) for Whale.

However, as our simulations suggest, these results will be sensitive to the degree by which the constant-rates assumption is violated. We note that the difference in estimated rates between WGDgc and Whale may already be an indication of considerable rate variation across branches of the species tree. Heterogeneity in the number of observed duplication and loss events across the species tree can be attributed to two sources: (1) genuine variation in duplication and loss rates across branches of the species tree or between major clades (i.e. variation in *λ* and *μ*), or (2) WGDs that are unaccounted for. It is possible in principle to estimate branch-specific duplication and loss rates, however such a model suffers from overparametrization and identifiability issues. Also convergence issues in the numerical optimization of the likelihood form a problem, as it becomes increasingly likely to get stuck in local optima for higher dimensional parameter spaces. This is especially true when WGD events are incorporated, because loss from non-retention and the birth-death process are confounded. Furthermore, we believe that an assumption of complete independence of duplication and loss rates across branches is also unreasonable from a biological perspective. These problems can be solved by assuming a model of rate heterogeneity and adopting a Bayesian approach, using the posterior distribution for retention rates or Maximum *a posteriori* (MAP) estimates for WGD inference. The benefit of such a Bayesian approach using informative priors is that we can overcome problems of statistical identifiability in the DL+WGD model and estimate branch-wise duplication and loss rates. Such an approach is however computationally very demanding. A more tractable, yet rather *ad hoc*, alternative is to perform ML estimation but with branches assigned to a limited set of rate classes, a so-called *local clock* model. Such an approach has been used for divergence time estimation using Maximum likelihood by Kishino and Hasegawa (1990), Rambaut and Bromham (1998) and Yoder and Yang (2000), and was particularly popularized in phylogenetics by the branch-tests for testing selection signatures in PAML (Yang 1998). We explore these approaches in the remainder of this paper.

First, we explore the effect of accounting for all previously described WGDs in the species tree of interest, as this might account for a considerable portion of the rate heterogeneity across these nine taxa. We performed the same analysis as for the angiosperm and seed plant WGD above but with six other WGDs that have been proposed along the species tree (Table 1). Note that we treated the so-called Gamma hexaploidy event as a WGD event (tetraploidy) and that we assumed only one WGD on the lineage leading to *O. sativa*. In both cases we expect a single WGD node marked in the species tree to capture most of the signal induced by the WGDs, but this obviously does not allow a clear interpretation of the resulting MLEs for the retention rates (but see further). MLEs for retention rates under the constant rates DL + WGD model are shown in Table 1. Most retention rates have an estimate considerably larger than 0, with only the angiosperm (*ϵ*) and eudicot (*γ*) events showing retention rates below 5%. For both *ϵ* and *γ*, the fit under the constant-rates model is significantly better with *q* equal to their respective MLEs (0.010 and 0.024 resp.) than with the corresponding rate set to 0. Such low retention rates however do not seem biologically relevant, and the better fit is likely due to some rate heterogeneity captured by the extra flexibility in the model coming from the WGD nodes. We note that accounting for these six additional putative WGDs resulted in a drop in the duplication rate MLE of over 30% 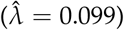, whereas the loss rate barely changed 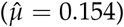. This is expected, as for an optimally chosen retention rate, some fraction of the duplications along a branch will be captured by this extra parameter, while the general incidence of loss should not be affected.

**Table 1:** Putative WGDs, their estimated dates and retention rate Maximum-likelihood estimates under different local-clock models. The constant rates model has one duplication and one loss rate for the whole tree. Local clock 1 estimates a duplication and loss rate for six rate classes (core angiosperms, *Amborella*, gymnosperms, *Sellaginalla*, bryophytes and all other branches). Local clock 2 estimates rates for the same rate classes as local clock 1, but with additional different rate classes for the stem branches of angiosperms, spermatophytes and tracheophytes.

| WGD | Mya | Date reference | Constant rates | Local clock 1 | Local clock 2 |
| --- | --- | --- | --- | --- | --- |
| Seed plants ($\zeta$) | 390 | Clark & Donoghue (2017) | 0.23 | 0.22 | 0.00 |
| Angiosperms ($\epsilon$) | 308 | Clark & Donoghue (2017) | 0.01 | 0.00 | 0.00 |
| *Picea abies* | 271 | Li *et al.* (2015) | 0.25 | 0.16 | 0.16 |
| Eudicots ($\gamma$) | 120 | Tiley *et al.* 2016 | 0.02 | 0.00 | 0.00 |
| *Oryza sativa* ($\rho$) | 91 | Clark & Donoghue (2018) | 0.25 | 0.21 | 0.21 |
| *Physcomitrella patens* | 66 | Vanneste *et al.* (2014) | 0.33 | 0.40 | 0.40 |
| *Arabidopsis thaliana* ($\beta$) | 55 | Vanneste *et al.* (2014) | 0.09 | 0.08 | 0.08 |
| *Arabidopsis thaliana* ($\alpha$) | 50 | Vanneste *et al.* (2014) | 0.16 | 0.16 | 0.16 |

Next we sought to account for some rate heterogeneity across lineages by assigning different branches to some fairly arbitrary rate classes. By doing this, we effectively relax our constant rates assumption to a constant rates assumption within each class. Again we mainly focus on the seed plant and angiosperm WGD, but will also direct our attention to retention rate estimates for the other six marked WGD nodes. Exploratory analyses indicated that a local clock with different rate classes for core angiosperms, gymnosperms, *Amborella, Sellaginalla*, bryophytes and all other branches seemed to explain a considerable amount of variation in rates while maintaining computational tractability (Figure S5, Supplementary Data 1). MLEs for the eight WGDs under this model are also shown in Table 1. Clearly, retention rates change in all directions compared to the constant rates model, with some retention rate estimates remaining relatively stable (*ζ, β* and *α*), some decreasing (*P. abies, O.sativa*) and one increasing (*P. patens*). Interestingly, the significant, but biologically implausible retention rates for the angiosperm and *γ* events are now estimated at 0, indicating that the retention rates indeed captured some rate heterogeneity in the constant rates analysis. Importantly, the seed plant WGD (*ζ*) has still a relatively high retention rate MLE.

Considering our focal WGDs, *ϵ* and *ζ*, it is important to note that the local clock 1 model discussed in the previous paragraph effectively assumes equal duplication and loss rates for the following branches: (a) the stem branch of tracheophytes, (b) the stem branch of spermatophytes and (c) the stem branch of angiosperms. We next checked the influence on the MLEs when relaxing this assumption (local clock 2, Figure S5, Table 1). This additional flexibility in the model results in the inability to detect a putative seed plant WGD. Several caveats are however to be considered. Most importantly, the model progressively becomes less identifiable the more independent rate parameters are allowed, and convergence issues for high dimensional problems make it increasingly likely to end up in a local maximum. We did however multiple runs and consistently obtained the values listed in Table 1. Secondly, under the local clock model, rates vary arbitrarily across rate classes, which may not be a biologically plausible model. For example, under local clock 2, the duplication-loss rate pairs (*λ, μ*) for the three successive stem branches (a, b & c) were estimated at (0.037,0.014), (0.464,0.239) and (0.140,0.276) respectively. Clearly, the three pairs are strongly different, and especially the duplication rate estimated for the stem branch of spermatophytes (b) is suspiciously high compared to the other estimates. These estimates would imply very large changes in gene duplication and loss rates on a relatively short phylogenetic time scale, which seems, although not impossible, highly implausible.

Both our simulations and investigations of the nine-taxon land plant data set using ML methods clearly illustrate the troubling influence of rate heterogeneity across lineages and associated assumptions when testing WGD hypotheses in a model-based framework. As already noted, an alternative solution to accommodate rate variation is to specify a prior distribution on duplication and loss rates across the species tree and adopt a Bayesian approach towards estimating parameters under the DL + WGD model. Besides omitting the need for specifying *ad hoc* rate classes, such an approach has the attractive property that we can employ models of duplication and loss rate evolution along the phylogeny, accounting for our *a priori* intuition that strong rate shifts should be relatively implausible. In the remainder of this study, we will explore such a Bayesian approach.

### Bayesian inference for the land plant data set

We applied a Bayesian approach for estimating branch specific duplication and loss rates and WGD retention rates to the nine-taxon land plant data set. As the MCMC algorithm is too computationally demanding for application on the full set of gene families, we restricted our analyses to a random subset of 1000 gene families. We note that our power simulations suggest that 1000 gene families should already provide sufficient power for biologically reasonable values of *q* (conditional on retention being more or less uniform over families). We tested the validity of our Bayesian approach on a limited set of simulations of 100 gene families for the nine-taxon tree from the previous section with six WGDs marked along the species tree (*α, β, γ, ϵ, ζ* and *P. patens*). The difference of the marginal posterior means for the sampled parameters from the simulated parameter values was centered around zero (Figure S6), suggesting that the marginal posterior means provide unbiased estimates for the parameters. WGD retention rates were well estimated for a set of 1000 gene families simulated under various strengths of rate heterogeneity for the same species tree (Figure 3). When the simulated rate variation was considerably stronger than the employed prior on the rate variability (*v* = 0.5 and *v* = 0.1 respectively), some duplication and loss rates have a strongly biased estimate, where the branch-wise estimate experiences shrinkage towards a common tree-wide rate. In general, these simulations show that a Bayesian approach using an informative prior on the variation in duplication and loss rates across the species tree can accommodate a considerable amount of rate heterogeneity and provides a means to estimate retention rates more reliably in the presence of rate variation.

**Figure 3:**
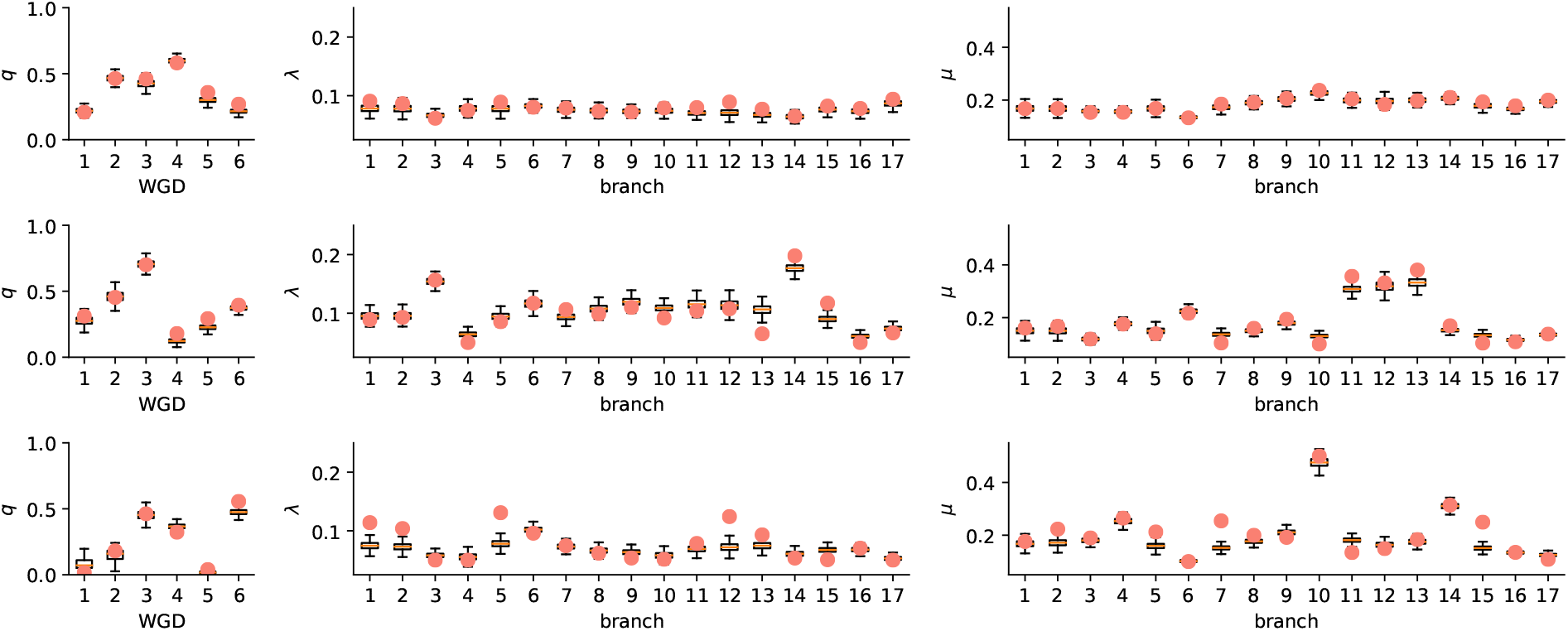
Bayesian posterior inference with Whale for simulated data sets. We simulated three data sets of 1000 gene families with duplication and loss rates sampled from the GBM prior with *v* = 0.10 (top row), *v* = 0.25 (middle row) and *v* = 0.50 (bottom row). We used a log-normal distribution 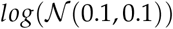 for the duplication rate at the root (*λ_τ_*) and log-normal distribution 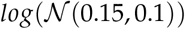 for the loss rate at the root (*μ_τ_*). We set the minimal duplication and loss rate to 0.05 and 0.1 respectively. Retention rates were sampled from a *Beta*(2,4) distribution and the geometric prior probability for the number of lineage at the root *η* was sampled from a *Beta*(10,1) distribution. We then performed Bayesian inference under the GBM prior with 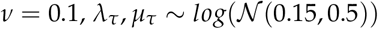, *q* ~ *Beta*(1,1) and *η* ~ *Beta*(4,2). The true (simulated) values are marked by dots, whereas the boxplots shows the sample from the posterior distribution obtained by MCMC with Whale.

Figure 4 (A) shows parameter estimates under the geometric Brownian motion (GBM) prior for duplication and loss rates and with 12 WGD hypotheses marked on the same nine-taxon tree (with *v* = 0.10). Among these 12 WGDs, we included two that are thought to be false (*S. moellendorffi* and *C. papaya* specific WGDs), five that are well-supported (*α, β, γ*, monocots and *P. patens*) and five that have been proposed with varying support (*ϵ, ζ*, gymnosperm, *G. biloba*, Pinaceae). Note that we only consider one WGD on the monocot branch here (but see below), whereas three are thought to have occurred in relatively close succession (*ρ, σ, τ*) (Jiao et al. 2014). For most of the WGDs that were also considered in our local-clock analyses we obtain qualitatively similar results. Using the approximated Savage-Dickey density ratio as a guide for hypothesis testing (see methods), we find support for *A. thaliana α* and *β*, no support for *γ*, strong support for a WGD event in *P. patens*, support for WGD in *O. sativa* and strong support for a seed plant event (*ζ*). A considerable drop in the retention rate for *P. patens* is observed, seemingly due to an elevated duplication rate relative to *M. polymorpha*. Interestingly, we find strong support for a WGD in the gymnosperm stem branch and no support for an event in *G. biloba*. Different from the local clock analyses using ML, is that we now do not find evidence for a Pinaceae specific WGD event, where the retention rate estimate under the local clock model was an artifact of the violated assumption of equal duplication and loss rates among gymnosperms. We again find no support for the *γ* WGD event nor for the hypothesized angiosperm specific WGD. Lastly, our analysis strongly rejects the non-hypothesized WGDs in *C. papaya* and *S. moellendorffii*, providing additional validation of our method.

**Figure 4:**
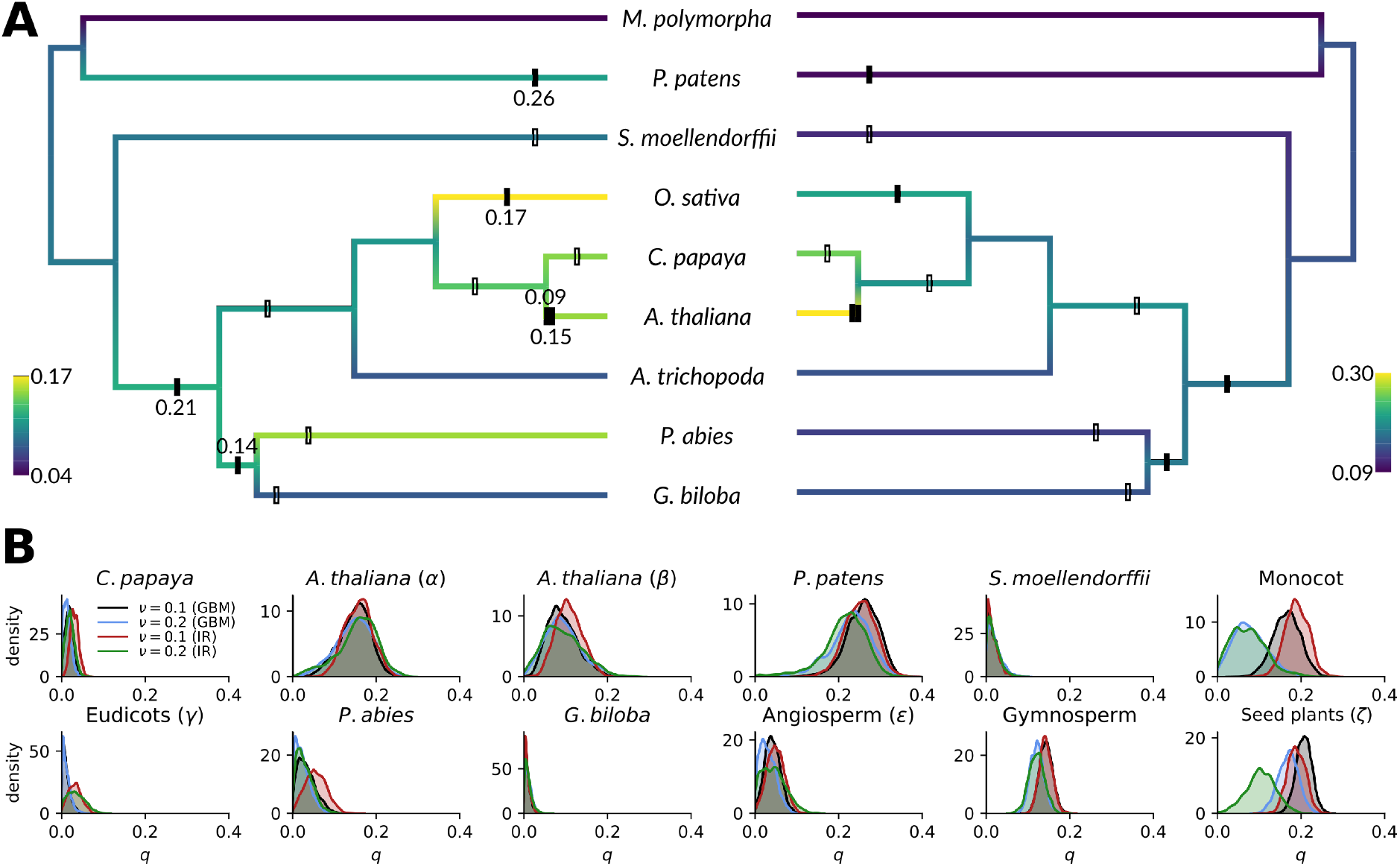
Bayesian inference of duplication, loss and retention rates under a geometric Brownian motion (GBM) (autocorrelated rates) prior for the nine-taxon angiosperm analysis. Inference is based on a random subset of 1000 gene families. (A) Posterior mean duplication (left) and loss (right) rates estimated under a GBM prior with *v* = 0.1 colored on the species tree lineages. Black bars indicate WGDs significantly different from zero and are annotated with the posterior mean for the relevant retention rate. The other bars indicate WGDs with a retention rate estimate not significantly different from zero. (B) Kernel density estimates for the marginal posterior distributions for the retention rates of all twelve WGDs. The different distributions show posteriors for different priors on the duplication and loss rate and different values of *v*.

We investigated the effect of using different prior assumptions by visualizing the marginal posterior densities for the retention rates acquired using different values of *v* and using an independent rates (IR) prior instead of the GBM prior (Figure 4 B). For most WGDs, using different prior assumptions has little influence on the marginal posterior distribution for the retention rate. A strong influence of the prior was observed for the hypothesized WGD on the monocot branch (*O. sativa*), where allowing more rate variability (larger *v*) resulted in a strong downward shift of the marginal posterior distribution for *q* and an increase of the estimated duplication rate for this branch for both the IR and GBM priors. A similar downward shift was observed for the putative seed plant WGD (*ζ*), however, here the shift was much stronger under the IR prior than the GBM prior. We observed that under the IR rates prior with *v* = 0.2, a biologically implausible loss rate was estimated for one of the branches stemming from the root (*μ* = 1.28), indicating that our prior parametrization was not strong enough to keep the parameter values within an *a priori* reasonable range (Figure S9). The results for the IR prior with *v* = 0.2 should therefore be interpreted with caution. We note that we obtained similar estimates for *η* in all analyses, ranging from 0.72 to 0.74. Together, our results for the nine-taxon analysis raise two important questions. (1) Why was the *γ* WGD in the eudicot ancestor not detected and (2) is there a gymnosperm specific WGD, seed plant specific WGD or both? We now turn to these questions in more detail.

In both the ML and Bayesian analysis, an anomalous result was obtained for the *γ* WGD event in the stem branch of eudicots. This event is well described, and is strongly supported by both *K_S_* distributions and synteny information, in particular from comparative analyses with *Vitis vinifera* (Jiao et al. 2012). We conducted a second analysis of a six-taxon data set including *V. vinifera* and *P. trichocarpa* in addition to the four angiosperms from the nine-taxon data set. In this data set we have two lineages that are thought to have not undergone subsequent WGD events after *γ* (*V. vinifera* and *C. papaya*), one lineage with one WGD event after *γ* (*P. trichocarpa*) and finally one lineage with two hypothesized WGDs after *γ* (*A. thaliana*). Curiously, in this analysis, support for a eudicot specific event is decisive, with a retention rate consistently estimated around 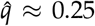 (Figure 5). A strong increase in estimated loss rates (*μ*) is observed in this six-taxon analysis compared to the nine-taxon results (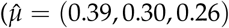 for the branches leading to the common ancestors of *A. thaliana - C. papaya, A. thaliana - P. trichocarpa* and *A. thaliana - V. vinifera* respectively, compared to 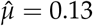 for the branch with the *γ* event in the nine-taxon analysis; both for the GBM prior with *v* = 0.1), and this may be the reason why the eudicot WGD was not detected in the nine-taxon analysis. It is unclear why such a difference in loss rate is observed between the two analyses. We further tested whether adding *P. trichocarpa* and *V vinifera* to the nine-taxon data set also resulted in a retention rate for *γ* significantly different from 0. Restricting our analysis to a smaller test set of 500 gene families (see methods) and only using the GBM prior with *v* = 0.1 for the sake of computational efficiency, we obtain a marginal posterior mean retention rate 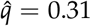 for the *γ* event (Figure S10). The loss rate estimates for the same three consecutive branches as noted above are 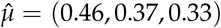. This suggests that the effect of taxon sampling is mainly due to breaking up long branches, not due to an overall different taxonomic breadth.

Further examining the results for the six-taxon data set, we again find support for the *α* and *β* events in the lineage leading to *A. thaliana*. Notably, we find a very high retention rate for *P. trichocarpa*, which is not unexpected, as genome analyses have revealed that the *P. trichocarpa* genome is for a very large part still present in duplicate (Tuskan et al. 2006). Support for three WGDs in close succession on the branch leading to *O. sativa* is limited, with posterior means for *q* between 0.08 and 0.1, and a Savage-Dickey density ratio favoring the model with *q* = 0 (1.0 < *K* < 1.2 in favor of *q* = 0 for all three WGDs under the GBM prior with *v* = 0.1). Changing the priors had different effects for the different WGDs, but overall did not result in different conclusions (Figure 5). For the *α* and *β* events in the *A. thaliana* lineage, increasing *v* mostly resulted in a wider distribution, whereas support decreased slightly for all WGDs in the monocot lineage. Both increasing *v* and using an IR prior had the effect of increasing the posterior mean for the *P. trichocarpa* WGD. The focal *γ* event was not affected by a change in the autocorrelation strength, whereas using an IR prior slightly decreased the posterior mean and increased the sensitivity to the *v* value. Overall, these results indicate strong support for the *α, β, γ* and Populus-specific WGD events and are suggestive, albeit not conclusive, for the hypothesized events in the monocot lineage. We note that a retention rate not significantly different from 0 does not ensure absence of WGD, but simply that the observed data can be explained by the duplication and loss rate. We also note that longer branches allow more flexibility in the duplication and loss rate under the GBM prior. A taxon sampling more focused on monocot species may therefore further constrain duplication and loss rates on internal branches and might be able to resolve the history of WGDs further. However this is beyond the scope of our current study and we refer to Jiao et al. (2014) for an in-depth study of WGDs in monocots. As for the nine-taxon data set, we obtain very similar *η* estimates in all analyses, ranging from 0.85 to 0.86.

**Figure 5:**
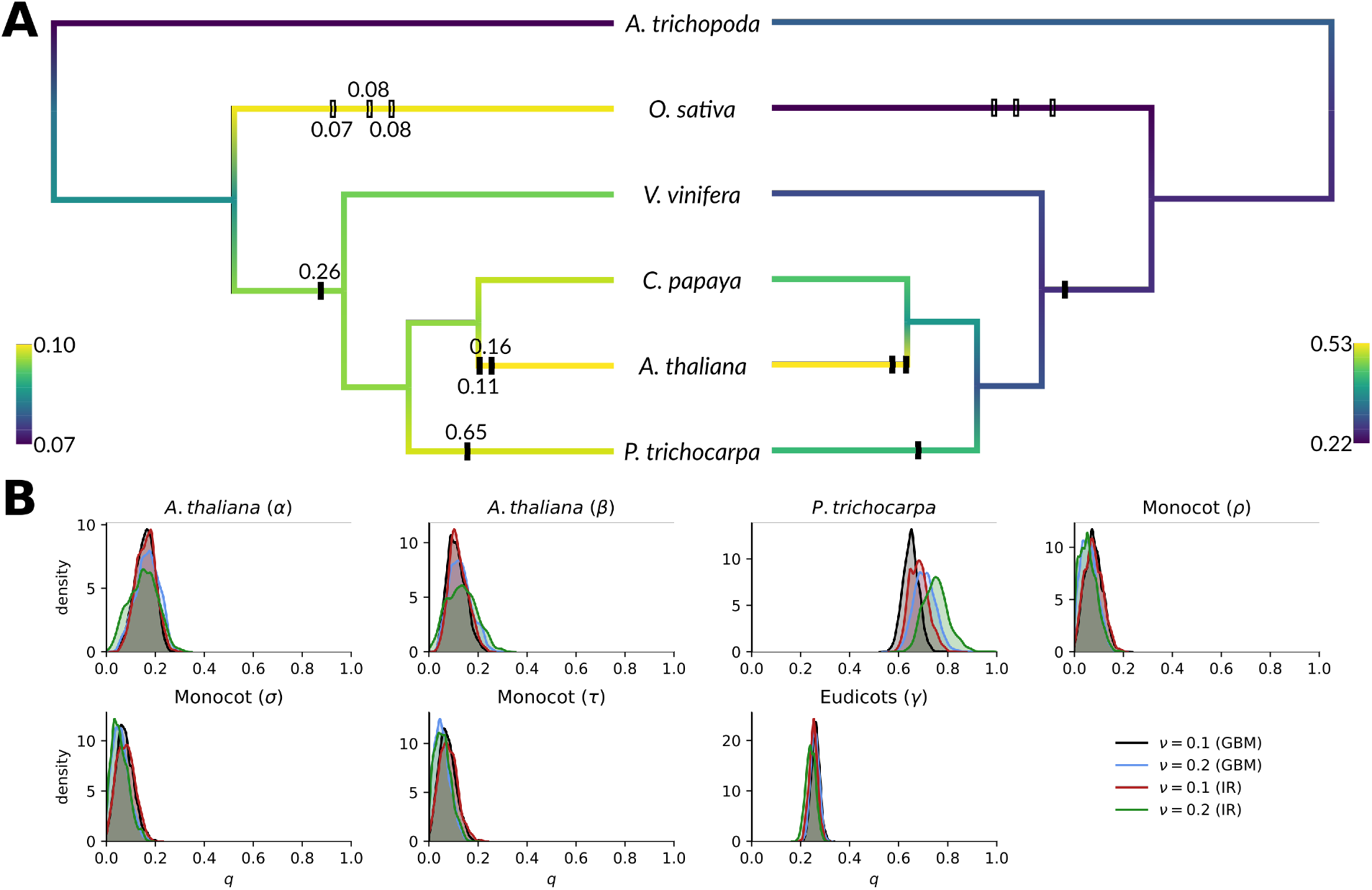
Bayesian inference of duplication, loss and retention rates under a geometric Brownian motion (autocorrelated rates) prior for the six-taxon angiosperm analysis. See Figure 4 for more information.

Our results for the nine-taxon data set demand a closer look at events in gymnosperms. Firstly, we did not find evidence for an event in the lineage leading to *P. abies* (Pinaceae), as was proposed in Li et al. (2015). Secondly, we found no evidence for an ancient WGD on the branch leading to *Ginkgo* around 300 Mya (before the divergence of cycads) but consistently found a non-zero retention rate for a hypothetical gymnosperm specific WGD event, which could explain the WGD signature in cycads and *Ginkgo*, one of the possible scenario’s suggested by Roodt et al. (2017). As the analysis for the *γ* WGD suggest that taxon sampling can exert an important influence on the analysis, we considered a five-taxon data set with four fully sequenced gymnosperm genomes, consisting of *G. biloba, P. abies, Pinus taeda, Gnetum gnemon* and *A. trichopoda*, to explore WGDs within gymnosperms in more detail. Remarkably, Bayesian inference under the Whale model found no support for any WGD events in gymnosperms (but see below) (Figure 6). Our analyses revealed *P. taeda* as a strong outlier, which forced us to fix the prior on the number of lineages at the root (*η* = 0.66) instead of sampling it during the MCMC. If we used the usual Beta hyperprior on *η*, this parameter was estimated at very low (biologically unreasonable) values, presumably to accommodate the outlier branch. Under the GBM model with *v* = 0.1 and *η* = 0.66, the mean duplication (loss) rate for the outlier branch was estimated at 0.51 (1.00) events per lineage per 100 My. The detection of such an outlier branch indicates that, despite our use of a fairly strong autocorrelation prior, strong deviating signals are nevertheless picked up for data sets of about 1000 gene families. This may have caused an upward bias in the rates nearby in the tree, however this is not obvious from the estimated rates. Inspection of some of the reconciled trees shows that often clades of very closely related genes are observed for *P. taeda*, which may just consist of alternative alleles erroneously assembled as separate genes and hints at a low assembly or annotation quality as the source of the outlier estimates. The absence of a Pinaceae specific event agrees with *K_S_* distributions for the two species (Figure 7). A suggestive low *K_S_* peak is present in some of the *K_S_* distributions of Pinaceae, which was interpreted as a shared Pinaceae WGD in Li et al. (2015). However, at least some Pinaceae do not show any peak (e.g. *Picea abies* and *Pinus taeda;* Figure 7), and other peaks in Li et al. (2015) are quite underwhelming, showing a signal that cannot be rightfully discerned from assembly or annotation artifacts due to heterozygosity for instance. We therefore believe that there is no convincing signal for a shared Pinaceae WGD, as was also originally concluded by Nystedt et al. (2013).

**Figure 6:**
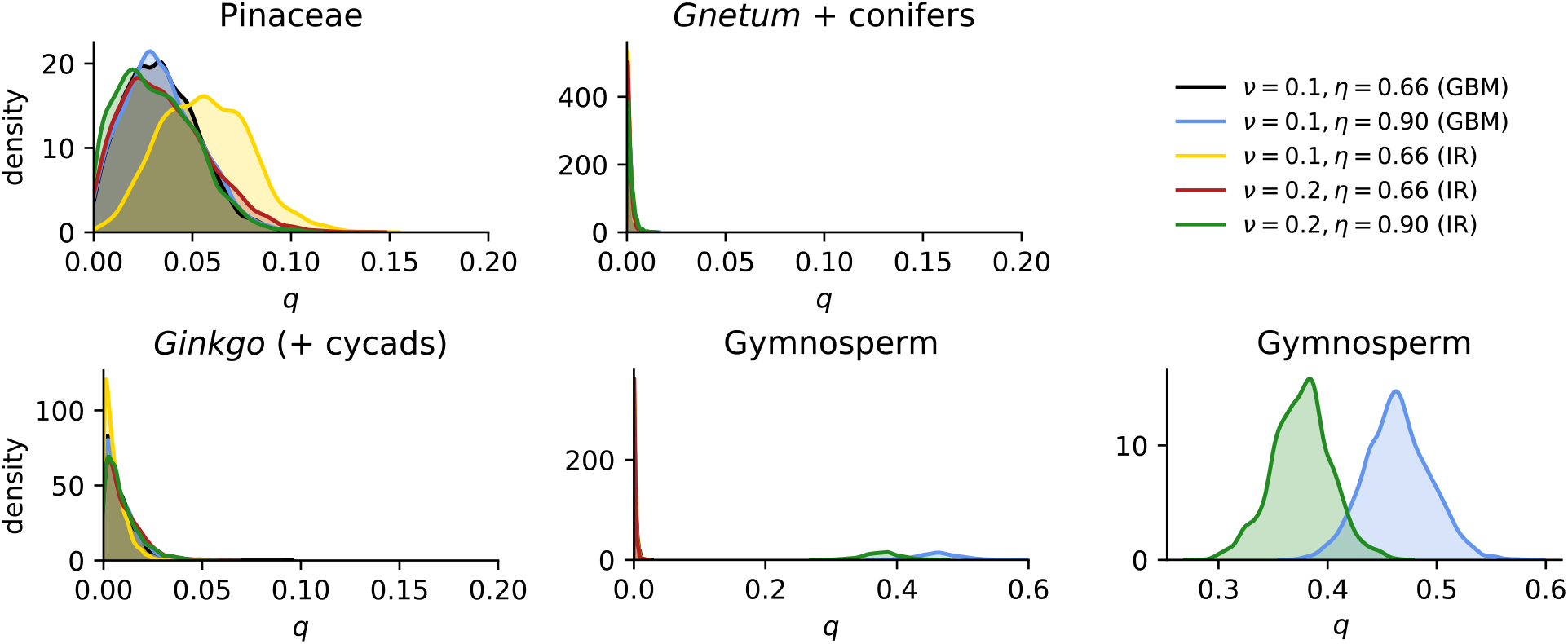
Bayesian inference of retention rates under different duplication and loss rate priors for the five-taxon analysis focusing on gymnosperms. The rightmost plot is a detail of the marginal posterior under the GBM prior with *η* = 0.9 for the hypothetical gymnosperm WGD event, which is also shown in the middle plot of the bottom row.

**Figure 7:**
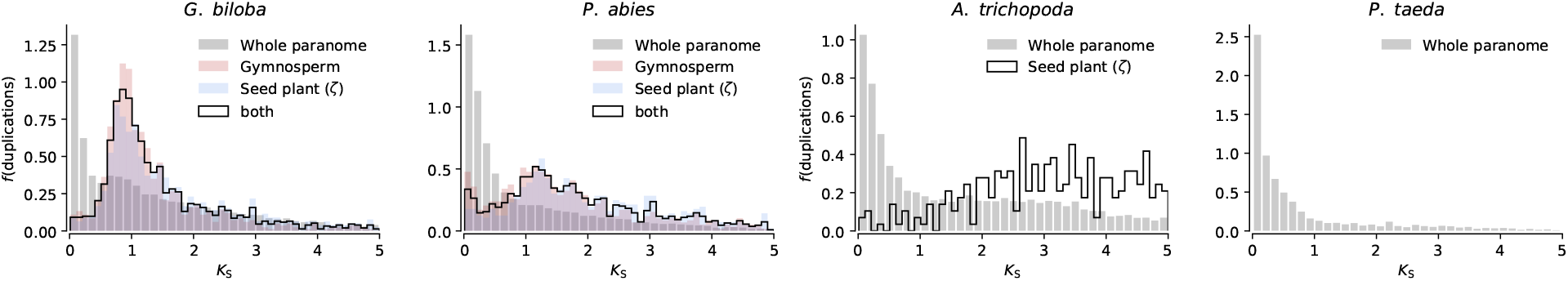
Node-averaged whole paranome *K_S_* distributions for *Ginkgo biloba, Picea abies, A. trichopoda* and *Pinus taeda*, with the *K_S_* distributions for those duplications reconciled to the hypothetical gymnosperm and seed plant WGD overlaid where appropriate. Note that the scale on the y-axis is a probability density, not a frequency or absolute number of duplications.

A possible explanation for the conflicting signals for the putative gymnosperm WGD in the nine-taxon and five-taxon analyses may be that it is an artifact due to a strong drop in duplication rate in the *Ginkgo* lineage compared to the gymnosperm stem and the lineage leading to *P. abies*. In the nine-taxon analysis, one can see that for the duplication rate on the gymnosperm stem branch, the average of the two daughter branches is close to the duplication rate estimated for the parental branch. In such a case, the autocorrelated rates model will strongly prefer a rate similar to the parental branch, and the extra flexibility from the WGD event is used to capture deviations thereof. Such an effect will be alleviated in the five taxon analysis due to the breaking up of the Pinaceae branch by including *Gnetum*. However, the fact that we obtained very similar duplication rate estimates for the gymnosperm stem branch under the IR prior (both *λ* ≈ 0.12) and that we obtain similar results in our restricted analysis of a 12-taxon data set which includes *Gnetum* (Figure S10) makes this explanation unlikely. Another possible explanation is gene tree uncertainty, causing the seed plant and gymnosperm events in the nine-taxon analysis to provide competing explanations for the same duplication events in a gene family. An example for this can be found in Figure S14, where different reconciled trees sampled from the posterior provide support for either a seed plant WGD, gymnosperm WGD or both. Lastly, uncertainty in the placement of the root may have caused conflicting results. Consider a duplication event in the gymnosperm stem with an outgroup gene from *Amborella*, both the scenario where the duplication is reconciled correctly in the stem of the gymnosperms and the scenario where the duplication is reconciled to the root, with one of the two lineages from the root being lost in *Amborella*, have a considerable probability *a priori*. As a result, specifying a WGD on a branch stemming from the root could result in less power to discern a WGD, and is very sensitive to the assumption on the number of lineages at the root. We tested this hypothesis by using a more stringent prior on the number of lineages at the root by specifying *η* = 0.9. Indeed, we observed a dramatic difference in the estimated retention rate for a putative gymnosperm WGD event (Figure 6). Assuming that the expected number of lineages at the root is ≈ 1.1 rather than 1.5, we can conclude that the five-taxon analysis supports a gymnosperm specific WGD. However it is unclear whether such an assumption is justified, especially since the presence of a putative seed plant WGD could cause a considerable number of orthogroups to actually contain two ancestors derived from the seed plant WGD. Together, the nine-taxon and five-taxon analyses suggest a gymnosperm specific WGD, however we must refrain from a final conclusion pending the availability of more and better gymnosperm genome assemblies. We reiterate that this illustrates how the assumptions on the number of lineages at the root can have a very strong effect when testing WGD hypotheses near the root of the species tree, a caveat which should also be important for the gene count method WGDgc of Rabier et al. (2014).

Our analyses provide evidence for a seed plant WGD, conflicting signals for an event in the stem of gymnosperms and no evidence for WGDs in gymnosperm lineages. A consequence of these findings is that the WGD signature observed in the *Ginkgo K_S_* distribution should trace back to the putative seed plant or hypothetical gymnosperm specific event. We sampled reconciled gene trees from the posterior distribution and extracted gene pairs in *Ginkgo* and *Picea* that are likely to have diverged after either the putative seed plant or gymnosperm WGD event (*i.e*. duplication nodes reconciled to the relevant WGD in at least 25% *(i.e*. on the order of 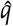) of the sampled trees), and examined their *K_S_* distribution (Figure 7). A strong peak in the distribution of these gene pairs is observed in *Ginkgo*, coinciding with the putative WGD signature in the whole paranome *K_S_* distribution. The signal in *Picea* is less clear, however it might be that the more dispersed peak observed here may also coincide with a signature present in the whole paranome distribution. Indeed, around *K_S_* ≈ 1, the *Picea* distribution shows a curious nick in the distribution. The distributions of the pairs reconciled to the gymnosperm and seed plant WGDs overlap completely, further suggesting that these may in fact correspond to a single event, with gene tree uncertainty obscuring the signal and diluting it over two consecutive branches. For *A. trichopoda*, only 288 of the putative seed plant WGD derived duplicates have an estimated *K_S_* < 5. We warn the reader that we show densities, not absolute numbers of duplications, on the y-axis in Figure 7, as the number of reliable duplication events extracted from the reconciled trees was way lower than the number of duplication events in the whole paranome *K_S_* distribution (Figure S11). This is most likely due to the orthogroup inference algorithm splitting up large and old orthogroups in multiple smaller orthogroups, whereas the whole paranome orthogroup clustering may result in larger clusters. Based on the combined results of our nine- and five-taxon analysis, we believe the most reasonable conclusion is that the hypothesized *ζ* WGD in the seed plant ancestor is likely to be true and that additional studies are required to evaluate the possibility of a gymnosperm specific WGD. Using different taxon samplings within gymnosperms might shed more light on this putative event, but high quality genome assemblies and complete annotations for gymnosperms remain scarce.

## Discussion

We developed a new model-based approach for evaluating WGD hypotheses in a phylogenetic context which jointly accounts for uncertainty in the gene tree and its reconciliation. Commonly used methods for inferring ancient WGDs using gene trees rely on a single maximum likelihood tree topology and its least common ancestor (LCA) reconciliation to a known species tree (e.g. Li et al. 2015; McKain et al. 2016; Zheng Li et al. 2018; Yang et al. 2018). Inference based on such an approach can however be misleading and induce biases that may lead to an inflation in the number of events inferred to explain the evolution of the gene tree in the species tree (Hahn 2007). To counter this, researchers have taken various measures, which in most cases amount to filtering out nodes with low bootstrap support when summarizing reconciliation results (Li et al. 2015; McKain et al. 2016; Zheng Li et al. 2018; Yang et al. 2018), but also heuristic branch swapping methods have been used (Li et al. 2016). Employing a probabilistic approach that considers the joint likelihood of a gene tree and its reconciliation alleviates the need for such *ad hoc* solutions to the problem of gene tree uncertainty. Indeed, Szöllősi, Rosikiewicz, et al. (2013) showed that joint inference using ALE resulted in strong decrease in the number of inferred events required to explain gene family evolution, and therefore, ironically, leads to more parsimonious explanations than parsimony-criterion based reconciliation of single maximum likelihood trees. Furthermore, a probabilistic model allows numerous possible modifications and improvements not considered in this study (e.g. accounting for missing data in transcriptomic data sets). We note that, besides being very efficient, the amalgamation approach has the merit that it only requires a sample from the posterior distribution over gene tree topologies. As a result, any software for MCMC-based phylogenetic tree inference can be used and the method we present here can thus readily make use of the vast array of models of sequence evolution that have been developed.

Whereas reconciliation based on parsimony criteria does not make explicit assumptions on duplication and loss rates, a probabilistic model necessarily does. We have shown through simulations and analyses of empirical data sets that statistical testing of presumed WGDs using gene counts or reconciled gene trees is strongly sensitive to these model assumptions. Our simulations show that the sensitivity of likelihood based tests under the constant rates DL+WGD model drops considerably in the presence of duplication and loss rate heterogeneity across the species tree, and the type I error rate rises unacceptably. Analysis of empirical data in an ML based setting with different local-clock models revealed that retention rate estimates can vary considerably when different assumptions on the duplication and loss rates are imposed. Accounting for rate heterogeneity across the species tree is therefore crucial to usefully employ model-based approaches.

These results also suggest that simulation based approaches to test for significant bursts in the number of duplications on particular branches in a species tree as used in (F.-W. Li et al. 2018; Zheng Li et al. 2018) should be interpreted with caution. Zheng Li et al. (2018) inferred reconciled gene trees using a ML tree and modified LCA reconciliation approach and summarized the number of duplications occurring on species tree branches of interest. To assess whether a particular number of duplicates corresponds to a significant increase in the number of duplications (possibly stemming from a WGD) they simulated gene tree topologies under the species tree of interest using a constant-rates DL model, with four sets of duplication and loss rates, which are estimated using gene count data. Statistical significance was assessed by comparing the number of duplicates in the DL model simulations with the actual data. We note that by using four sets of rates in their simulations, the authors account partly for rate heterogeneity across gene families, but not species tree lineages, whereas our simulations suggest however that the latter has a more dramatic effect. While such an approach can of course be valuable and indicative of WGD events, one should be aware that the null hypothesis that is actually tested here is whether the number of duplications on a species tree branch inferred using the particular reconciliation approach is approximately equal to the number inferred for trees generated under a constant-rates linear birth-death process operating along the species tree phylogeny. The relationship of such a hypothesis with ‘significant bursts in the number of duplications’ is by no means obvious, in part because of rate heterogeneity across the tree, but also because of possible methodological biases in heuristic reconciliation approaches. Of course, we do not claim that such simulations are meaningless, but merely question the interpretation of the statistical test based thereupon.

Problems with rate heterogeneity in model-based testing of WGD hypotheses using the gene count method implemented in WGDgc were implicitly addressed by Tiley et al. (2016) by performing analyses on reduced four-taxon species trees to complement their global analysis. Such an approach, however, still constrains the outgroup and ingroups for a particular WGD event to have the same duplication and loss rates, which is a particularly problematic assumption for very ancient WGDs. For example, as we show here, the duplication rate in mosses or *Sellaginella* is considerably lower than for example the duplication rate in *Arabidopsis*, and this would compromise analyses where we use for example *P. patens* as outgroup to assess the putative seed plant WGD. Specifying local-clock models therefore provides an alternative to these reduced taxon-set analyses that can overcome such limitations. However, both approaches suffer from the same possible arbitrariness in either taxon sampling or rate class assignment. Nevertheless, we believe careful analysis using ML estimation with local-clock models can provide valuable insights for relatively small species trees.

Incorporating models of rate heterogeneity as prior information in a Bayesian approach provides a promising but challenging solution. Employing these priors allows us to estimate branch-wise duplication and loss rates without having to assume complete independence of rates across the species tree, and effectively serves as a way to perform regularization by shrinking branch-wise estimates to a common tree-wide rate. Here we used both an independent rates (IR) prior and autocorrelated rates (GBM) prior, where the latter takes into account a correlation structure imposed by the species phylogeny whereas the former does not. Our analyses using this Bayesian approach suggest that several putative WGD events that are commonly accepted might need revision. In particular, our results for the hypothesized angiosperm specific WGD event are in line with recent doubts cast upon the original analyses that lead to the proposal of this event (Ruprecht et al. 2017). We believe that evidence for an angiosperm specific WGD event is insufficient, and that the general acceptance of this event is unjustified. We obviously do not rule out the possibility of an angiosperm specific WGD, this is something that our methods cannot rule out with certainty, but simply state that stronger evidence is required. Importantly, our study of the *γ* WGD indicated that breaking up a long branch with a hypothesized WGD by employing a more dense taxon sampling can result in considerably different retention rate and gene duplication and loss rate estimates. The same problem may affect our ability to detect a hypothetical angiosperm WGD event. Sadly, however, we cannot further break up the stem branch of angiosperms, *Amborella* being the first extant diverging lineage known after the divergence of gymnosperms. This fundamental issue provides a challenge for proving whether or not a WGD preceded the divergence of angiosperms by phylogenetic means.

Several other proposed WGD events were also not supported by the present study, such as the proposed Pinaceae WGD (Li et al. 2015) or *Ginkgo* specific WGD (Guan et al. 2016), and we suggest that, pending evidence from high-quality genome assemblies, these hypotheses should not be regarded as well-supported. Whereas our results can be sensitive to our prior assumptions on the duplication and loss rates, the effects are case-dependent and most of our results do not change qualitatively when employing different priors. We believe prior settings that resulted in our ability to detect well-supported WGDs (e.g. *A. thaliana α* and *β, P. patens, P. trichocarpa*, eudicot *γ*, monocot WGDs) and give negative results where analysis from other sources indicate absence of WGD (e.g. *C. papaya, S. moellendorffi*) can be regarded as reasonable. Additionaly, our detailed analysis of the *γ* WGD event and hypothetical gymnosperm WGDs showed that the method is still sensitive to taxon sampling, and that this is not only so for the autocorrelated rates prior. Lastly, Whale supports the hypothesized seed plant WGD (*ζ*), and detected a gymnosperm specific WGD signal in our nine-taxon data set. Further analyses of a data set focusing on gymnosperms provided conflicting results on the hypothetical gymnosperm specific WGD, and leave the possibility that the signal might reflect an artifact. Our analyses must therefore necessarily remain inconclusive. We believe the most reasonable interpretation in the light of previous research and currently available data and methods is a seed plant specific WGD event, and that additional analyses might shed light on the possibility of a WGD event in the gymnosperm ancestor. However, we stress that all possible WGD hypotheses on these ancient branches should remain to be entertained.

Several critical assumptions of the Whale method in its current implementation should be noted. Firstly, as all other approaches, Whale relies on accurately inferred orthogroups and alignments, and essentially assumes that these represent the true homology relationships across a set of genomes. If large and old orthogroups are erroneously split up in multiple distinct subfamilies, our ability to detect ancient events will be reduced. The comparison of *K_S_* distributions derived from our reconciliations to the whole paranome distributions suggests this might play a role (Figure S11). Our method assumes that gene families evolve independently but with the same rates. This assumption may be relaxed by incorporating a model of rate variation across families on top of the rate variation across species tree branches. This is effectively analogous to the Gamma models of rate heterogeneity across sites in likelihood and distance based phylogenetic tree inference (Yang 1994). However, simulations show that across family heterogeneity has little influence on the estimated retention rates, and we believe that the possible better model fit might be of little added value considered in the light of the additional computational time needed. We also assume that different gene lineages evolving within a species tree branch have identical duplication and loss rates, as the duplication and loss rate is a property of a species lineage in our model. Lastly, the model for WGDs adapted from Rabier et al. (2014) makes multiple convenient yet possibly problematic assumptions. This model assumes that retention is independent across families, whereas it is known that gene families belonging to different functional categories show different patterns of gene retention following WGD, which is thought to stem from dosage effects (Maere et al. 2005; Tasdighian et al. 2017). It is unclear whether such effects would however influence our rate estimates. Perhaps more importantly, our model assumes that the process of diploidization, where a polyploid genome is resolved back to a diploid genome (with associated gene loss captured by our retention rate parameter), occurs in a time frame that is very short compared to the time scale of the species tree. This assumption may be problematic for autopolyploidy events on relatively short internal branches of the species tree. For such branches, diploidization of some gene families may only be complete after the speciation event following the WGD, resulting in a pattern that was called Lineage specific ohnologue resolution (LORe) by Robertson et al. (2017). More generally, violation of this assumption may lead to incomplete lineage sorting of duplicated genes. Relatedly, our model does not discriminate between allo- and autopolyploidy events. This means that for allopolyploidy events, the model assumes that the two donor species diverged within the species tree branch on which the WGD is hypothesized, and the WGD node effectively corresponds to this divergence event. If this assumption is suspected to be violated, meaning that there is a sister species tree branch which is more closely related to one of the donor genomes, our model does not hold and one should resort to a parsimony-based method that can account for allopolyploidy (Thomas et al. 2017). Accounting for these complexities in a probabilistic framework is another challenge for future research, and would require more sophisticated models that explicitly model the polyploid phase of the lineage under consideration.

It is also important to consider that the linear birth-death process might be an overly simple model of gene family evolution. In particular, our model does not account for incomplete lineage sorting, and incorporating the multi-species coalescent in our framework to account for the possibility of deep coalescence would be an interesting future development. Furthermore, we note that while a staggering number of models of sequence evolution exist, models of gene family evolution have been very limited. Indeed, all previous research efforts on model-based gene tree reconciliation have modeled gene family evolution as a linear birth-death process (Arvestad et al. 2009; Rasmussen and Kellis 2011; Szöllősi et al. 2012; Szöllősi, Rosikiewicz, et al. 2013; Rabier et al. 2014). Posterior-predictive model checking shows that while the DL+WGD model seems to fit the observed data reasonably well, several interesting things should be noted that could inform improved models of gene family evolution (Figure S16). The most important insight is that real data sets show an overrepresentation of single-copy core gene families (*i.e*. families with exactly one gene for each species). This indicates that models with a notion of ‘essentiality’, where the transition from one gene copy to extinction of a gene family in a species tree lineage has a different rate than the transition from *n* to *n* − 1 gene copies (with *n* > 1), could provide an improved fit of the data. Similarly, better models to account for duplication and loss rate variation across the tree can obviously benefit our approach. We believe these might be fruitful further research directions. However, any attempt to introduce more complex models in the Bayesian setting would require more efficient computation. Indeed, we had to restrict most of our Bayesian analyses to reduced sets of gene families (~1000) and limited sets of species (~10) for the sake of computational feasibility.

To conclude, we stipulate the crucial interaction between duplication and loss rates and putative WGD hypotheses that forms the core message and problem of our present study. On the one hand, any strong deviation from the assumptions on the duplication and loss rates may result in the retention rate capturing some signal not accounted for by the used model of rate heterogeneity, with the resulting possibility of false positive inference. On the other hand, too much flexibility in the model (*i.e*. few assumptions on duplication and loss rates), will make it more plausible that the signal of a WGD is absorbed in the duplication and loss rate estimate, which is allowed to vary freely in a biologically implausible way. In our view, a Bayesian approach using a prior on the rates is more reasonable than either a constant rates or local-clock assumption. In order to obtain sufficient efficiency and reasonable rate estimates we were required to set the crucial parameter *v*, governing the amount of rate variation in both the autocorrelated (GBM) and uncorrelated (IR) rates priors, to a fixed value. However, increasing this parameter within a reasonable range had minor effects on our results. Therefore we believe that the assumptions made in our study are justified in light of the methods currently at our disposal. Additionally, we showed that taxon sampling remains important in a model based approach, and that interactions of WGD hypotheses with uncertainty on the gene tree root can provide misleading results. Improved phylogenetic models of both gene family evolution and polyploidization, as well as better and more efficient methods for Bayesian inference under the Whale or similar models, may provide further insights and corroborate or contradict the findings we presented. Finally, our analyses of WGDs across the land plant phylogeny showed that some of the hypothesized WGD events are not beyond reasonable doubt, and we believe that our understanding of genome evolution would benefit from continuous critical re-examination of the many proposed events in the light of new methods and data.

## Methods

### Amalgamated likelihood estimation with a duplication, loss and WGD model

#### Amalgamated likelihood estimation (ALE)

We use the ALE method (Szöllősi, Rosikiewicz, et al. 2013) to efficiently estimate the joint gene tree - reconciliation likelihood under a known dated species tree *S*(*i.e*. where divergence times are assumed to be known), a model of sequence evolution 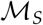 and a model of gene family evolution 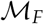 for a multiple sequence alignment *D*. The ALE approach allows approximating the likelihood

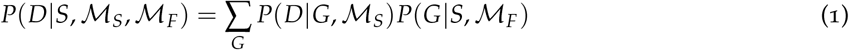

where *G* denotes the gene tree topology and the sum is over all possible gene tree topologies for the sequences in D. This approximation is achieved efficiently by performing amalgamation (Szöllősi, Rosikiewicz, et al. 2013) using the conditional clade distribution (CCD) (Höhna and Drummond 2012; Larget 2013), which can be derived from a sample from the posterior distribution of tree topologies using any Bayesian phylogenetic inference program. Under the principle of conditional independence of subtrees, Höhna and Drummond (2012) and Larget (2013) showed that conditional clade probabilities (CCPs) provide an accurate approximation of the posterior probability of a gene tree topology. The conditional clade probability of a clade *γ* in tree *G* with daughter clades *γ*′ and *γ*″ is defined recursively by

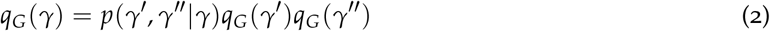

where *p*(*γ*′,*γ*″|*γ*) is computed from the CCD simply as the fraction *f*(*γ*′,*γ*″)/*f*(*γ*), where *f* denotes the frequency of the clade (or pair of clades) observed in a sample from the posterior distribution of tree topologies under inferred under 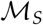. If *γ* is a leaf in *G*, we have *q_G_*(*γ*) = 1, ending the recursion. Szöllősi, Rosikiewicz, et al. (2013) showed that given the marginal split frequencies (*i.e*. *p*(*γ*′,*γ*″|*γ*) for all *γ* observed in the sample), the CCD is the maximum entropy distribution for the sample space of all tree topologies. Under the principle of conditional independence of subtrees, *q_G_*(Γ) now approximates 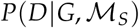, where Γ denotes the *ubiquitous* clade, *i.e*. the clade encompassing all members of the gene family under consideration. Szöllősi, Rosikiewicz, et al. (2013) used this result to design a set of recursions amenable to dynamic programming to approximate the joint gene tree - reconciliation likelihood for all gene trees that can be *amalgamated* from the observed clades in the MCMC sample. If we, following the notation of Szöllősi, Rosikiewicz, et al. (2013), denote the probability under 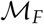 of observing the subtree rooted in node *u* of the gene tree in branch *e* of the species tree at time *t* as *P_e_*(*u, t*), and analogously Π_*e*_(*γ, t*) for a clade *γ*, the ALE approximation amounts to

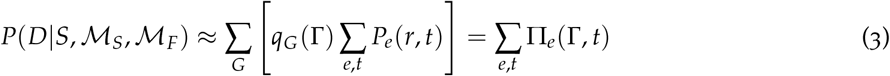

Where *r* denotes the (unknown) root of the gene tree. For more details on the ALE approach, we refer to Szöllősi, Rosikiewicz, et al. (2013). Below, we elaborate on our adaptations of (3) and the main recursions for Σ_*e,t*_ Π(Γ, *t*) to accommodate a different model of gene family evolution including duplication, loss and WGDs (DL + WGD). We also discuss further restrictions on the likelihood computation by accounting for filtering procedures and alternative assumptions on the gene tree root.

#### Probabilistic gene tree reconciliation using a DL + WGD model

We implemented the ALE method using a model of gene family evolution accounting for duplication, loss and whole genome duplications (WGDs) first proposed by Rabier et al. (2014). A simple linear birth-death model with birth (duplication) rate *λ* and death (loss) rate *μ* is used to model duplication and loss, whereas for every WGD node marked in the species tree an extra parameter *q*, the retention rate, is included to model massive loss after WGD. Every lineage that passes through the WGD node is forced to duplicate, with a duplicate copy retained with probability *q* and lost with probability 1 − *q*. This effectively models the number of retained WGD-derived duplication nodes for a particular WGD across a set of gene families as a binomially distributed discrete random variable with parameter *q* and sample size the number of extant gene lineages just before the WGD event. We note that this is a very crude model for the phylogenetic consequences of a WGD, which assumes that rediploidization occurs in a very short time frame compared to the evolutionary timescales studied.

We adapt the recursions of Szöllősi, Rosikiewicz, et al. (2013) to implement the DL + WGD model with the ALE method for approximating the joint tree - reconciliation likelihood. Following Szöllősi, Rosikiewicz, et al. (2013), we discretize each branch of the species tree *S* into *d* segments of length Δ*t* and define *t* to increment from the present to the root (i.e. *t* = 0 at the leaves of *S*). Using this ‘sliced’ species tree *S*, we compute for every clade *γ* at every time point *t* along *S* the probability of observing *γ* at time *t* for every branch *e* of *S* in terms of its daughter clades *γ*′ and *γ*″ observed in the sample, and we denote this probability as Π_*e*_(*γ, t*). We define Π_*e*_(*γ*,0) ≡ 1 if *e* is a branch leading to a leaf of *S* and *γ* corresponds to a gene of the corresponding species, and 0 otherwise. The within-branch recursion, allowing for duplication and simple propagation within a slice of length Δ*t*, is given by

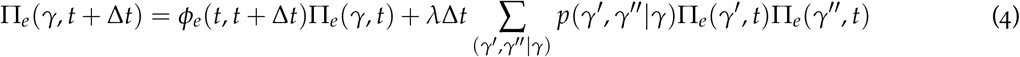

Where *ϕ_e_*(*t, t* + Δ*t*) denotes the probability of propagation of a single lineage, conditioning on the lineage being non-extinct, which can be computed from properties of the linear birth-death process (see Appendix). The first term accounts for the possibility of no event with observed descendants in time interval [*t, t* + Δ*t*], whereas the second term accounts for the possibility of duplication at rate *λ* in the time slice, with both lineages origination from the duplication event having observed descendants. This recursion is effectively a stripped down version of the one found in Szöllősi, Rosikiewicz, et al. (2013), as we do not consider horizontal gene transfer events here.

At a speciation node at time *t*, the incoming branch *e* gives rise to two child branches *f* and *g*. The speciation event can be observed in the gene tree, leading to a split of a clade *γ* in two daughter clades *γ*′ and *γ*″. Alternatively, the speciation event may not be observed in the sample due to a loss event following the speciation event, with a probability given by the extinction probability at the trailing edge of the child branch where the loss occurred *ϵ_g_*(*t*). Accommodating both scenario’s, the recursion at speciation nodes becomes

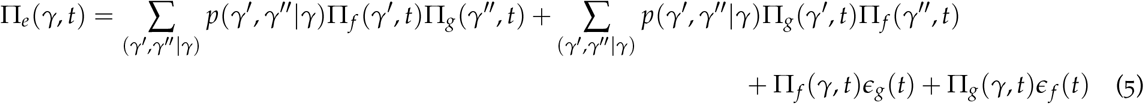

Where the first two terms account for observed speciation events and the last two terms for speciation and loss events. This is identical to the expression in Szöllősi, Rosikiewicz, et al. (2013). The extinction probability *ϵ_e_(t)* is defined as the probability that a gene at time *t* in branch *e* leaves no observed descendants. These probabilities can be computed analytically using properties of the linear birth-death process using a postorder traversal along the species tree as in Arvestad et al. (2009), Rasmussen and Kellis (2011) and Rabier et al. (2014) (see Appendix).

At WGD nodes, we employ the WGD model of Rabier et al. (2014). However, we consider all possible reconciliations at the WGD node, in a way similar to the within-branch and speciation node recursions, as opposed to Rabier et al. (2014), who only considered most parsimonious reconciliations. Both lineages from the WGD derived duplication can either be retained with probability *q*, or one lineage may be lost through massive gene loss post-WGD with probability 1 − *q*. We also have to consider the cases where one of the lineages from a retained WGD node is not observed due to extinction in branches below the WGD node. Summing over these three possible scenario’s, we have for a WGD node at time *t_j_* in branch *e*

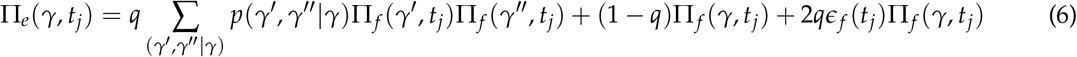

Where *f* is the branch below the WGD node. This recursion is illustrated in Figure S12.

The recursions outlined in this section enable the calculation of the joint gene tree - reconciliation likelihood under the DL + WGD model given a sliced species tree *S* and a sample from the gene tree topology posterior using dynamic programming. We next consider the special considerations needed when handling the root of *S* and the ubiquitous clade Γ.

#### Prior on the number of lineages at the root and conditioning

Szöllősi, Rosikiewicz, et al. (2013) compute the overall joint likelihood as 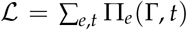, effectively summing over *all* possible ways the gene tree might be reconciled to the species tree. This allows *de novo* birth of gene families on branches below the root without a probabilistic model thereof (but note that an ‘origination’ probability can be included as in Szöllősi et al. (2012) to account for this). Multiple lineages at the root (*r*) of the species tree are allowed in this model by adding a virtual branch above the root where duplication events can happen. Here we do not consider the possibility of origination in arbitrary subtrees of *S* by filtering gene families such that they contain at least one gene in both clades stemming from the root of *S*. Instead of adding the virtual branch above the root, we follow Rabier et al. (2014) and Csűrös and Miklós (2009), and specify a geometric prior distribution on the number of lineages at the root. Under the geometric prior with success probability *η*, the probability of observing *n_r_* = *s* lineages at the root with *s* ≥ 1 is given by

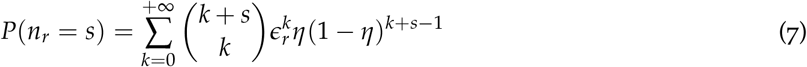

where *ϵ_r_* = *ϵ_f_*(*τ*)*ϵ_g_*(*τ*) with *f* and *g* denoting the two branches stemming from the root and *τ* denoting the age of the root (i.e. the species tree height). This expression accounts for the possibility of lineages at the root that are unobserved due to extinction in *S*, which occurs with probability *ϵ_r_*. Under this prior, the expected number of lineages at the root is simply 1/*η*. The above expression can be rephrased as

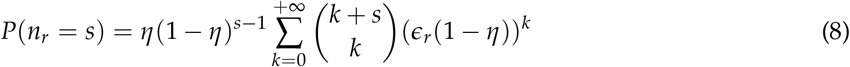

and since |*ϵ_r_*(1 − *η*)| < 1, the above expression converges to

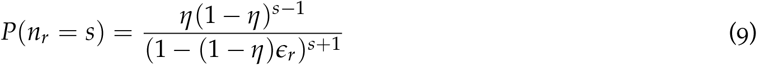

For *s* lineages reconciled at the root of the species tree with subtree probabilities *P_r_*(*T_i_,τ*); *i* ∈ {1,⋯,*s*}, the probability of observing the gene tree *T* given the species tree and a geometric prior with parameter *η* is

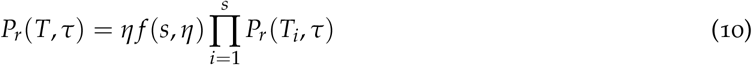

where we have defined the function *f*(*s, η*) as

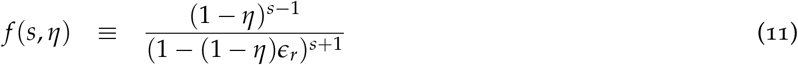

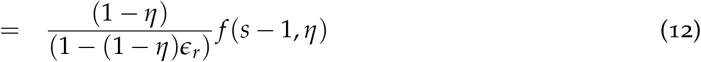

This recursive formulation allows a recursion for the probability Π_*r*_ (*γ, τ*) of observing a clade *γ* reconciled to the root. If we denote 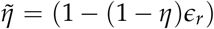, we can express Π_*r*_(*γ, τ*) in terms of its daughter clades *γ*′ and *γ*″ as

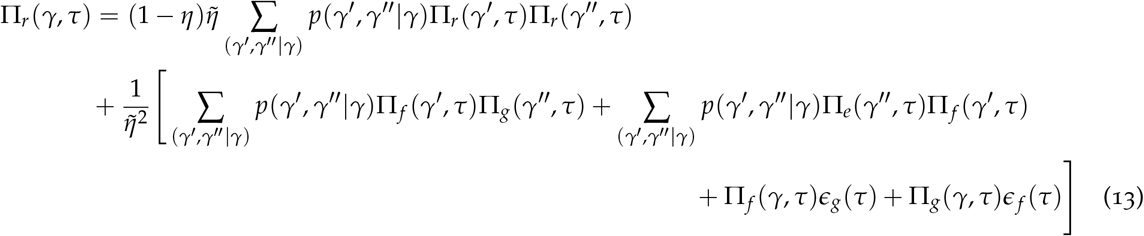

Figure S13 illustrates this recursion at the root for two given reconciliations. Under this model, the total likelihood is given by *η*Π_*r*_(Γ, *τ*) instead of 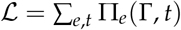. For a single gene family *D*, we have instead of (3) above

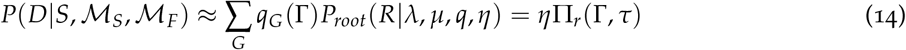

Rabier et al. (2014) further showed that conditioning on the filtering step described above as well as the fact that the gene family is not extinct is crucial for obtaining unbiased estimates for *λ* and *μ*. This conditioning step can be performed after computation of the likelihood as in Rabier et al. (2014), and does not require adaptation of the specified recursions. Throughout our study, we condition on the gene family having at least one member in each clade stemming from the root. If we denote the unconditional likelihood for family *i* by *P*(*D_i_*), we have *P*(*D_i_*) = *P*(*D_i_, i* observed) = *P*(*D_i_|i* observed)*P*(*i* observed). For our filtering procedures the conditional likelihood is than given by

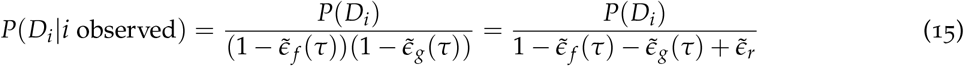

where 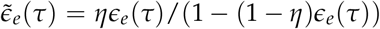 denotes the extinction probability at the root time *τ* in branch *e* under the geometric prior distribution (Rabier et al. 2014). After computing the three-dimensional reconciliation matrix, reconciled trees can be sampled by stochastic backtracking along the sum as in ALE (Szöllősi, Rosikiewicz, et al. 2013).

### Inference by Maximum likelihood and MCMC

We implemented the Whale method described above in the Julia programming language. Evaluating the likelihood for a representative set of 1000 gene families from the nine-taxon data set (see below) takes ~ 3.5s on a single 2.6 GHz CPU. We use the Nelder-Mead downhill simplex method to numerically maximize the log likelihood under the DL and DL + WGD model as a function of the duplication, loss and retention rate parameters and acquire the MLEs thereof. In order to assure computational feasibility, we implemented the likelihood evaluation routine in parallel to harness modern multi-core computing infrastructure. The implementation allows arbitrary local clock models, allowing anything in between a constant duplication and loss rates model and branch-wise rates model.

To accommodate rate variation in a model-based framework, we implemented an adaptive Metropolis-within-Gibbs Markov chain Monte Carlo algorithm (MCMC) to sample from the posterior distribution under different priors on the duplication, loss and retention rates. We implemented a geometric Brownian motion (GBM) prior on the duplication and loss rates (Thorne et al. 1998; Rannala and Yang 2007) to model correlated rates across the species tree. Under this model of rate drift, given the rate *r_A_* at the parental node, the logarithm of the rate *r* at some node different from the root follows a normal distribution 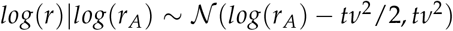, where *t* is the branch length between the node and its parent, and *v* is inversely proportional to the autocorrelation strength. We used the implementation of Rannala and Yang (2007), which is an adaptation of the approach in Thorne et al. (1998), where rates are estimated for the midpoints of the species tree branches. In this model we use a lognormal prior on the duplication and loss rate at the root, where we set the mean to 0.15 and scale to 0.5 throughout this study.

We also implemented an independent rates (IR) prior, similar to Rannala and Yang (2007), where all duplication and loss rates are modeled as independent and identically distributed random variables from the same log-normal distributions with means *λ*_0_ and *μ*_0_ respectively and fixed standard deviation *v*. Note that *v* has a different interpretation in the IR model compared to the GBM model. Nevertheless, in both models *v* is proportional to the amount of rate variation tolerated by the model. In both the GBM and IR models, we also employed a hyperprior on *v*. However, in settings with data sets of a size relevant for this study (>50 gene families), large *v* values are strongly preferred despite specifying a strong prior on *v* which favors values close to 0. When this happens, rates can vary unrestrictedly across branches of the species tree, which can lead to identifiability issues and is unreasonable from a biological point of view. This problem stems from having highly informative data for a biologically unreasonable model for which we currently have no proper alternative. As a result, we used a fixed *v* value in both models, and evaluate the sensitivity of the resulting posterior distributions on this parameter by running different chains with different *v* values. In all models we use a Beta prior *Beta*(1,1) for the retention rates (*i.e*. a uniform distribution in the interval [0,1]) and a *Beta(4*,2) prior on the success probability *η* of the geometric distribution for the number of genes at the root of the species tree. We choose a *Beta*(4,2) prior because it has a mean of 0.66 (which we use as fixed *η* in the ML setting) but has more mass on values of *η* > 0.5 than for example a *Beta*(2,1) prior (which has the same mean), corresponding with our prior beliefs that there should be only one or a few lineages at the root if we have proper orthogroups.

We implemented a Markov chain to sample from the posterior distribution for duplication and loss rates (*λ* and *μ* resp.), retention rates (*q*) and the expected number of lineages at the root (1/*η*) given by

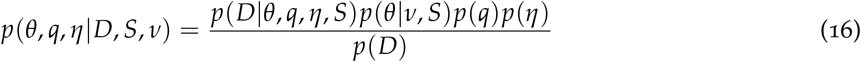

Where *θ* denotes the vector of branch-specific duplication and loss rate pairs (*λ_i_, μ_i_*) and *q* the vector of retention rates. We used the following update cycle for each MCMC iteration (where *x|y* indicates that we update the parameter (vector) *x* conditioning on the current *y*):

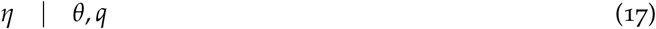

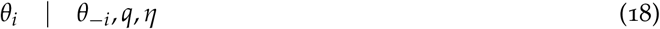

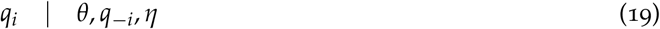

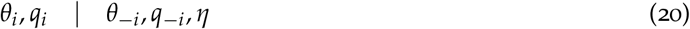

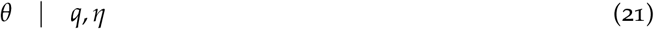

where (19) and (20) are only performed for branches with WGD nodes. The subscript *i* indexes branches, and indicates that this update is for each branch separately, and the subscript −*i* denotes the vector of all parameter values for branches excluding index *i*. All updates use a symmetric random walk proposal with standard deviation subject to vanishing adaptation using the methods described in Roberts and Rosenthal (2009). In brief, for every parameter we measure the acceptance rate in batches of 50 MCMC iterations and adapt the variance of the proposal kernel for that parameter based on the monitored acceptance rate. If the acceptance rate is below the target value, the logarithm of the variance of the proposal kernel is increased by a factor *δ*, whereas it is decreased when the acceptance rate exceeds the target value. To ensure that the resulting Markov chain has the desired stationary distribution, *δ* is a decreasing function of the batch number, approaching 0 asymptotically. This entails that the magnitude of adaptation diminishes as the MCMC chain runs, which assures that the chain retains the desired stationary distribution. Convergence of the chain was monitored using trace and autocorrelation plots, and we computed the effective sample size (ESS) for every parameter (Brooks et al. 2011). In general we found that for our problems a burn-in of about 1000 generations followed by 10,000 generations sufficed to achieve a good sample from the posterior, with the minimal ESS ≈ 100, and most parameters having an ESS > 500.

WGD hypotheses are assessed by inspection of the marginal posterior distribution for the associated retention rate parameter. Additionally, Bayes factors can be computed from the marginal posterior distributions by means of the Savage-Dickey density ratio. Let us denote by *p*(*ψ, q*) the prior density under the model with a particular WGD of interest (representing the alternative hypothesis) and *p*_0_(*ψ*) the prior density under the model without the focal WGD (representing the null hypothesis), where *ψ* stands for all parameters different from the retention rate of the WGD of interest. Since *p*(*ψ|q* = 0) = *p*_0_(*ψ*), a result from Dickey (1971) and Verdinelli and Wasserman (1995) shows that the Bayes factor can be computed from the marginal posterior density at *q* = 0, *i.e*.

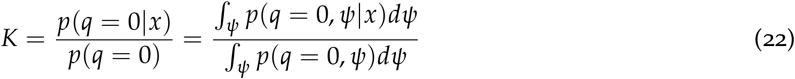

where *x* denotes the data. We approximate the numerator in this expression by the value at *q* = 0 of the boundary-corrected kernel density estimate of the marginal posterior distribution for the retention rate of interest. Posterior predictive model checks were done by simulating gene families from the posterior and comparing gene family sizes and gene counts for each species with the observed gene family data.

### Data analysis and software availability

Amino acid sequence data was downloaded from PLAZA 4.0 (Van Bel et al. 2017) for all species except *G. biloba* and *P. taeda*, where data was gathered from the TreeGenes (Wegrzyn et al. 2008) database. Orthogroups were inferred with OrthoFinder (version 2.1.2 Emms and Kelly 2015) under default settings. Orthogroups that did not contain at least one gene in both clades stemming from the root (i.e. at least one gene in mosses and one gene in tracheophytes) were discarded, to rule out the possibility of *de novo* origin of orthogroups in the species tree under study. We also filtered out the 5% largest gene families for the sake of computational efficiency (10% for the 12-taxon analysis in Figure S10). For every orthogroup, an amino acid alignment was inferred using PRANK (Löytynoja and Goldman 2008) with default settings and a sample from the posterior distribution was obtained by means of MCMC using MrBayes 3.2.6 (Ronquist et al. 2012) under the LG+Γ4 model. After a burn-in of 10,000 generations, the chain was sampled every 10 generations over a total of 100,000 generations for every family. The conditional clade distribution (CCD) was inferred from the resulting posterior sample using ALEobserve from the ALE software suite. For the 12-taxon analysis we further filtered out orthogroups that had a very ‘large’ CCD *(i.e*. a very high number of clades) exceeding 500 diferent clades. Throughout, we use a dated species tree derived from the study of Morris et al. (2018) for reconciliation purposes, with branch lengths in units of 100 million years. *K_S_* distributions for *G. biloba, Picea abies, Pinus taeda* and *Amborella trichopoda* were computed using the wgd package using default settings (Zwaenepoel and Van de Peer 2019). The Whale program is implemented in Julia and available from https://github.com/arzwa/Whale.jl. A nextflow pipeline for all preparatory computation (alignment, tree inference and CCD construction) as used in this study is available from https://github.com/arzwa/Whaleprep.

## Acknowledgements & Funding

We thank Wandrille Duchemin and Gergely Szöllősi for some initial help with regard to technical details of the ALE approach. This work was supported by the European Union Seventh Framework Programme (FP7/2007-2013) under European Research Council Advanced Grant Agreement 322739—DOUBLEUP [to Y.V.d.P]; and a PhD Fellowship of the Research Foundation—Flanders (FWO) [to A.Z.].

## Appendix

### Calculation of extinction probabilities

Recall that we have discretized the species tree *S* in *d time slices* of length Δ*t* for each branch of the species tree (Figure S17). Time runs from the present *t* = 0 to the root *t* = *τ*. We will now compute for each time slice the probability that a lineage entering the slice leaves no observed descendants, and we denote this probability as *ϵ_e_*(*t*) for time *t* on branch *e*. We first note the following features of the problem occuring at the extremities of the branches in *S*:

a. *ϵ_e_*(0) = 1 if *e* is a branch leading to a leaf in *S*.
b. For a branch *e* speciating at time *t* into two daughter branches *f* and *g* we have *ϵ_e_*(*t*) = *ϵ_f_*(*t*)*ϵ_g_*(*t*)
c. For a WGD node with retention rate *q* at time *t* that marks the boundary between branch *e* and branch *f* we have *ϵ_e_*(*t*) = *qϵ_f_*(*t*)^2^ + (1 − *q*)*ϵ_f_*(*t*)

To compute the extinction probabilities we will need the transition probabilities from the linear birth-death process. The transition probability *P_n|a_*(*t*) for *a* lineages to undergo duplication at rate *λ* and loss at rate *μ* such that there are *n* lineages after time *t* is given by

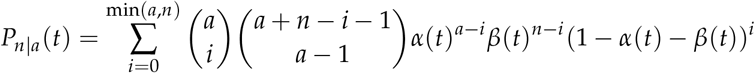

with

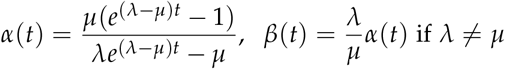

and

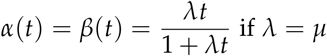

(see for example Bailey 1964). Here *α*(*t*) is the probability of a single lineage going extinct over time *t*.

We now show how to compute extinction probabilities over the times slices (each of length Δ*t*) of a branch recursively. Consider a branch *e* in *S* leading to a leaf of *S*. By (a) above, we have at time *t*_0_ = 0 that *ϵ_e_*(*t*_0_) = 0. At time *t*_1_ = *t*_0_ + Δ*t* = Δ*t* in the time slice above, we have the extinction probability given by *ϵ_e_*(*t*_1_) = *α*(Δ*t*). If we go another time slice higher up (to *t*_2_) in the tree we do not only consider the probability of extinction within the time slice *α*(Δ*t*) but also the possibility of any other birth-and death scenario giving rise to *i* lineages at *t*_1_ below, followed by extinction of the *i* lineages each with probability *ϵ_e_*(*t*_1_), that is

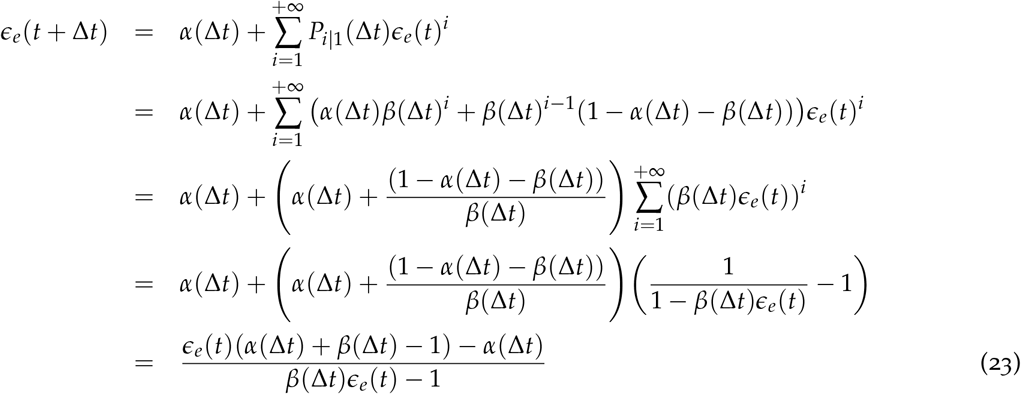

In combination with conditions (a), (b) and (c) above, this expression allows to calculate the extinction probabilities at any time point along the discretized species tree *S* using a postorder traversal of *S*.

### Calculation of the single gene propagation probabilities

We will now derive the recursions for the propagation probabilities *ϕ*(*t, t*′). The propagation probability is the probability that a single lineage entering a time slice at time *t* ‘propagates’ through the time slice to generate exactly one lineage at the end of the time slice (time *t*′) which has observed descendants at the present (*t*_0_ = 0). Consider again a branch *e* in *S* leading to a leaf of *S*. At time *t*_0_ = 0 we have *ϕ*(*t*_0_, *t*_0_) = 1, since this corresponds to an observed lineage. At time *t*_1_ = *t*_0_ + Δ*t* = Δ*t* we simply have *ϕ*(*t*_0_, *t*_0_ + Δ*t*) = *P*_1|1_(Δ*t*). At *t*_2_ = *t*_1_ + Δ*t*, we consider the possibility that the single lineage undergoes an arbitrary number of duplications and losses such that there are *i* lineages at *t*_1_, of which *i* − 1 go extinct in the time slice below. We therefore have

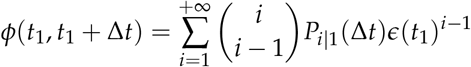

Which is very similar as for the extinction probabilities outlined above. Note that 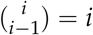. If we work this out for the general case, we get

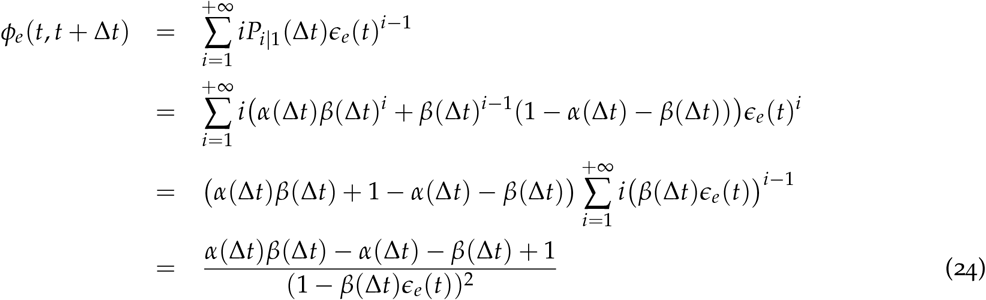

These recursions allow to compute the single gene propagation probabilities given *λ μ*, and the extinction probabilities that can be computed using the recursions from the previous section.

## Supplementary figures

**Figure S1:**
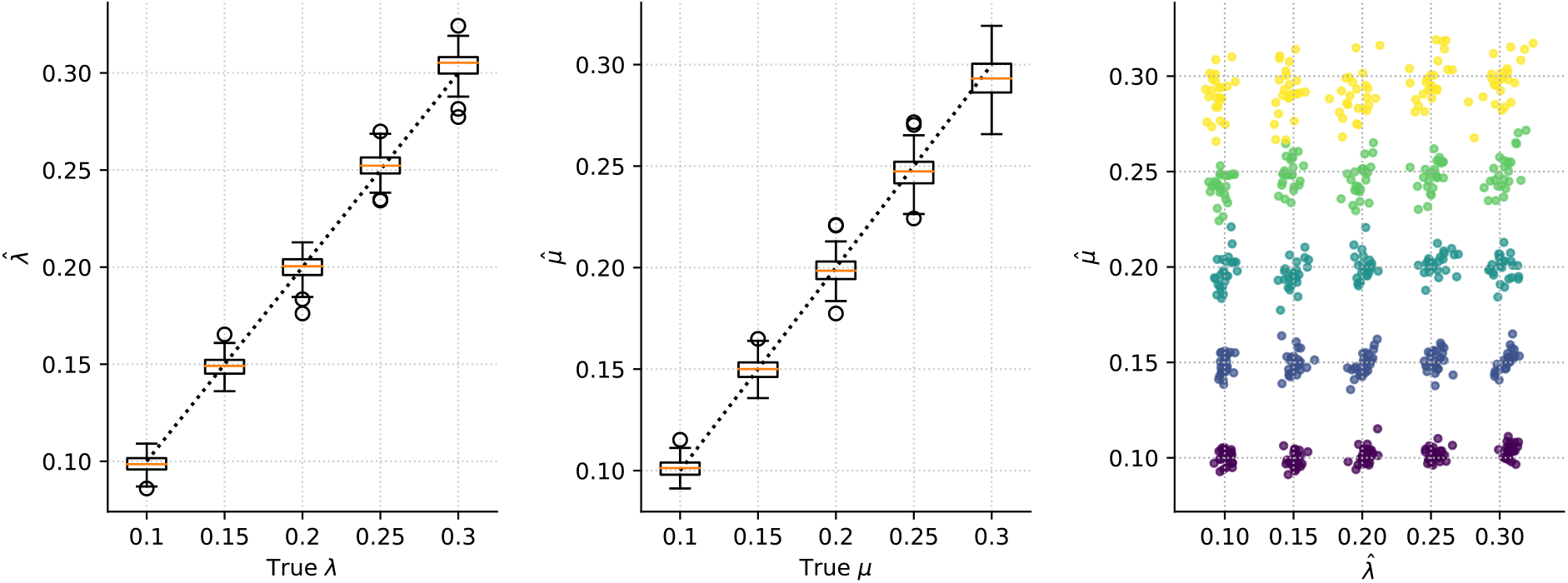
MLEs for the duplication 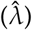 and loss 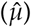 rates for 25 data sets of 200 simulated trees for every *λ, μ* pair. The dashed line marks the simulated values.

**Figure S2:**
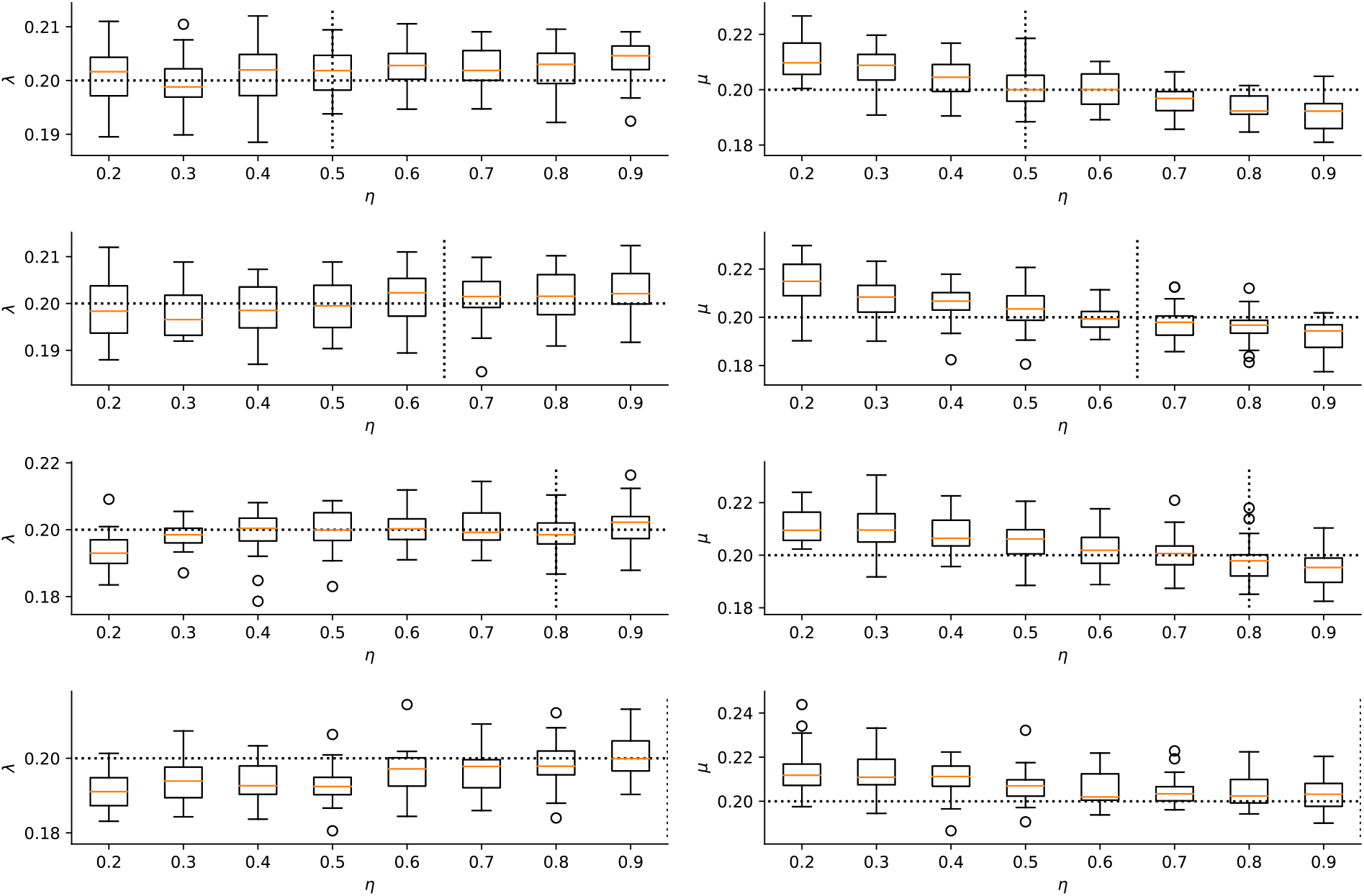
Duplication and loss rate estimates under different simulated and used parameterizations of the prior on the number of genes at the root. The x-axis shows the value used for *η* in the ML estimation of the rates whereas the simulated *η* value is marked by the vertical dashed line. Every box-plot is based on 25 replicates.

**Figure S3:**
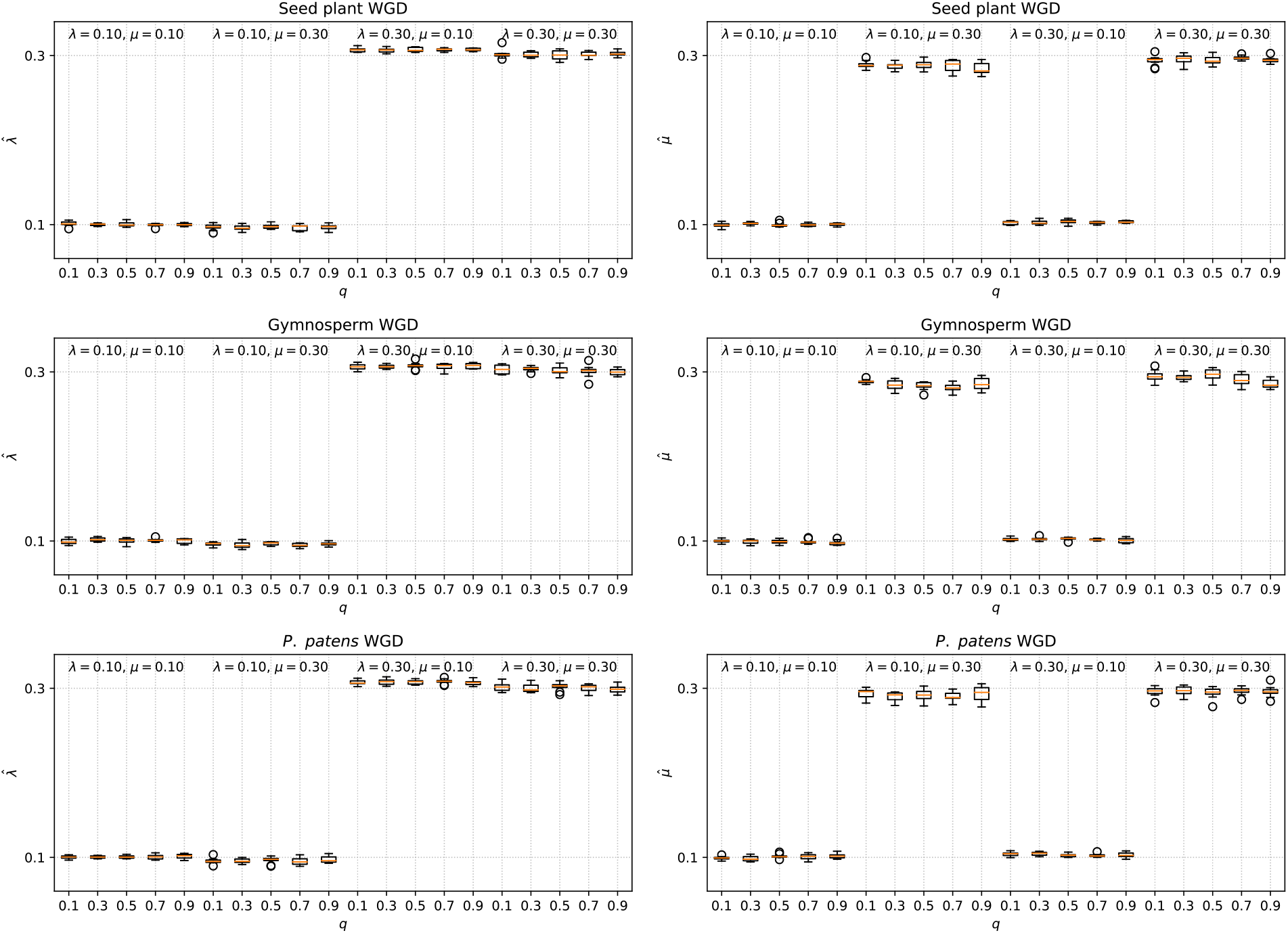
MLEs for the duplication and loss rate for different simulated WGD scenario’s and duplication and loss rates. Simulations of 10 times 500 gene families were done for a 10-taxon tree with constant duplication and loss rates across the tree. Each box-plot is based on 10 replicates for data sets of 500 gene families. The species trees used for the simulations are shown in Figure 1.

**Figure S4:**
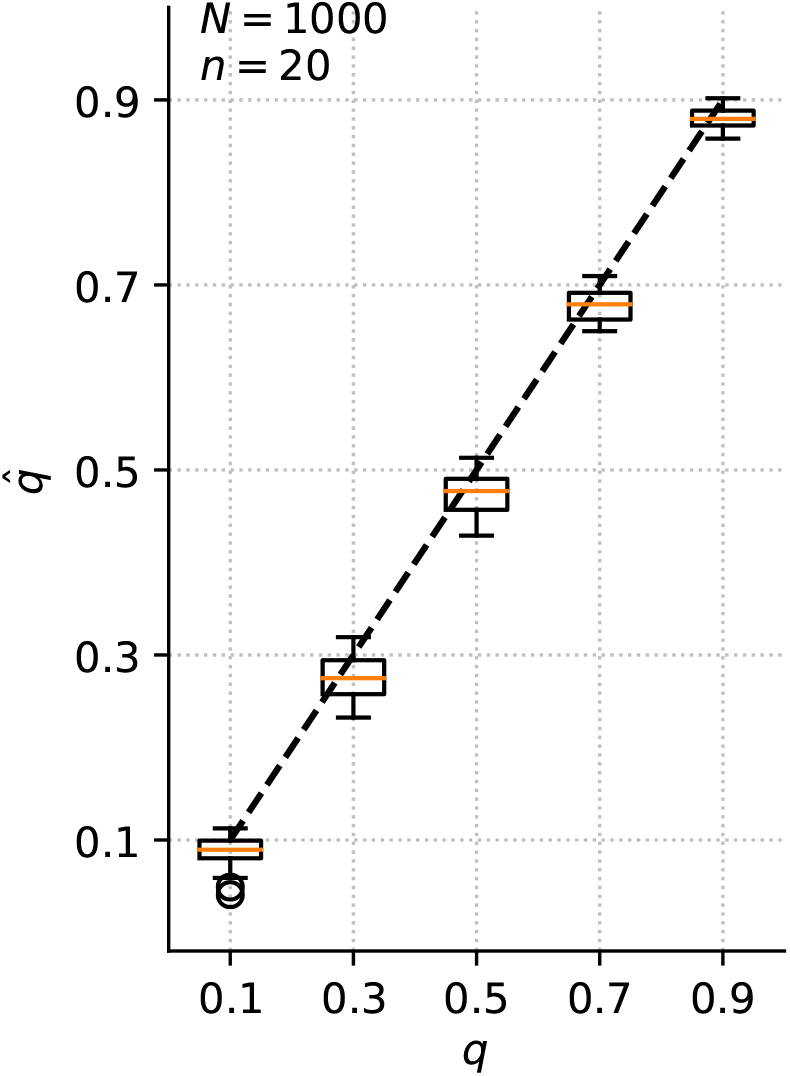
MLEs for the retention rate 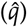 based on 20 simulations of 1000 gene trees where duplication and loss rates for different families were drawn independently from a Gamma distribution with shape *k* = 2 and scale *θ* = 0.1.

**Figure S5:**
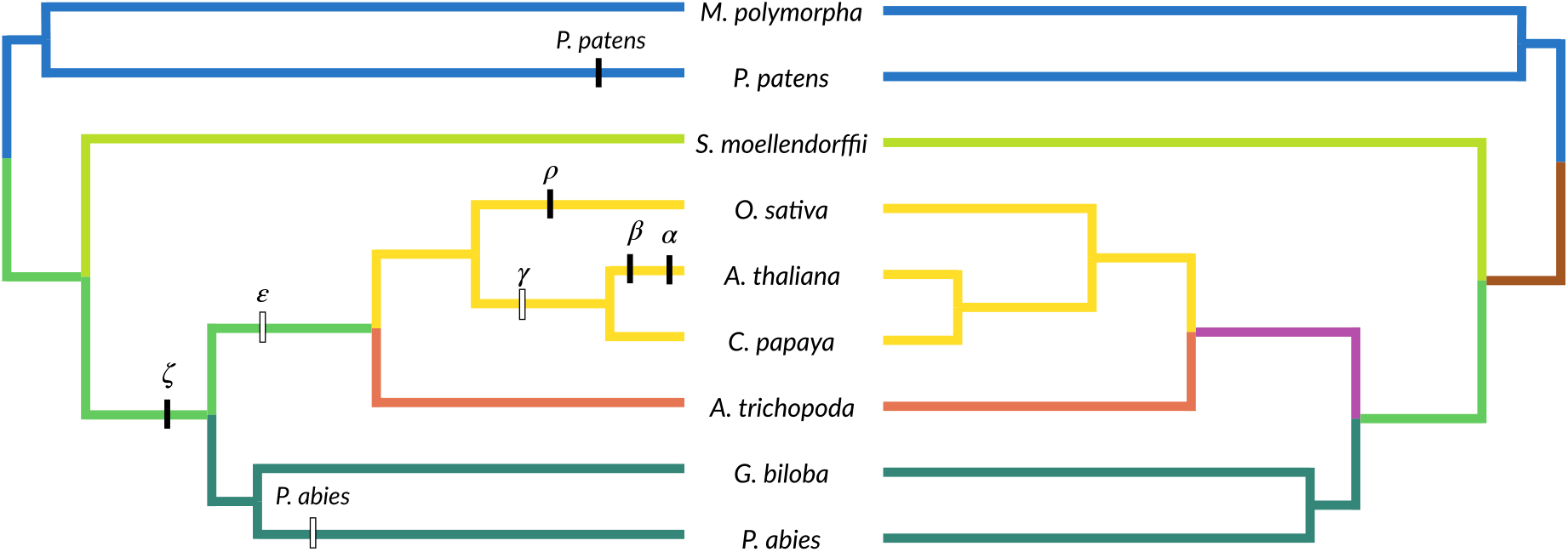
The eight WGDs investigated and the local-clock models used in the maximum likelihood inference for the nine-taxon data set. Different colors mark different rate classes. The tree on the left shows the rate classes for local-clock 1 whereas the right tree shows the rate classes for local-clock 2.

**Figure S6:**
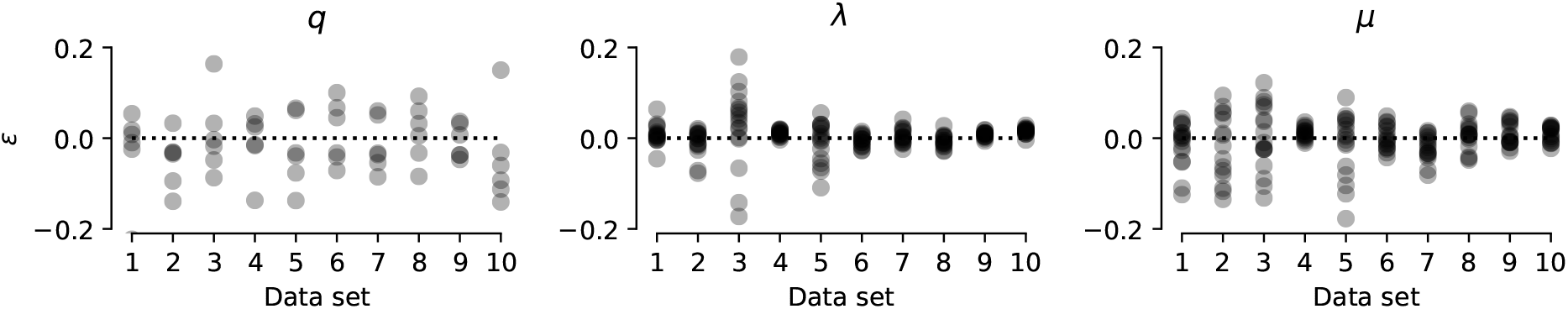
Error (*ϵ*) of the posterior means for Bayesian inference with Whale. We simulated 10 data sets of 100 gene families with duplication and loss rates sampled from the GBM prior (*v* = 0.1) with a log-normal distribution 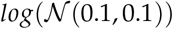 for the duplication rate at the root (*λ_τ_*) and log-normal distribution 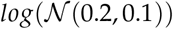 for the loss rate at the root (*μ_τ_*). Retention rates were sampled from a *Beta*(2,4) distribution and the geometric prior probability for the number of lineage at the root *η* was sampled from a *Beta*(10,1) distribution. We then performed Bayesian inference under the GBM prior with *v* = 0.1, 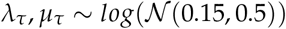, *q ~ Beta*(1,1) and *η ~ Beta*(4,2).

**Figure S7:**
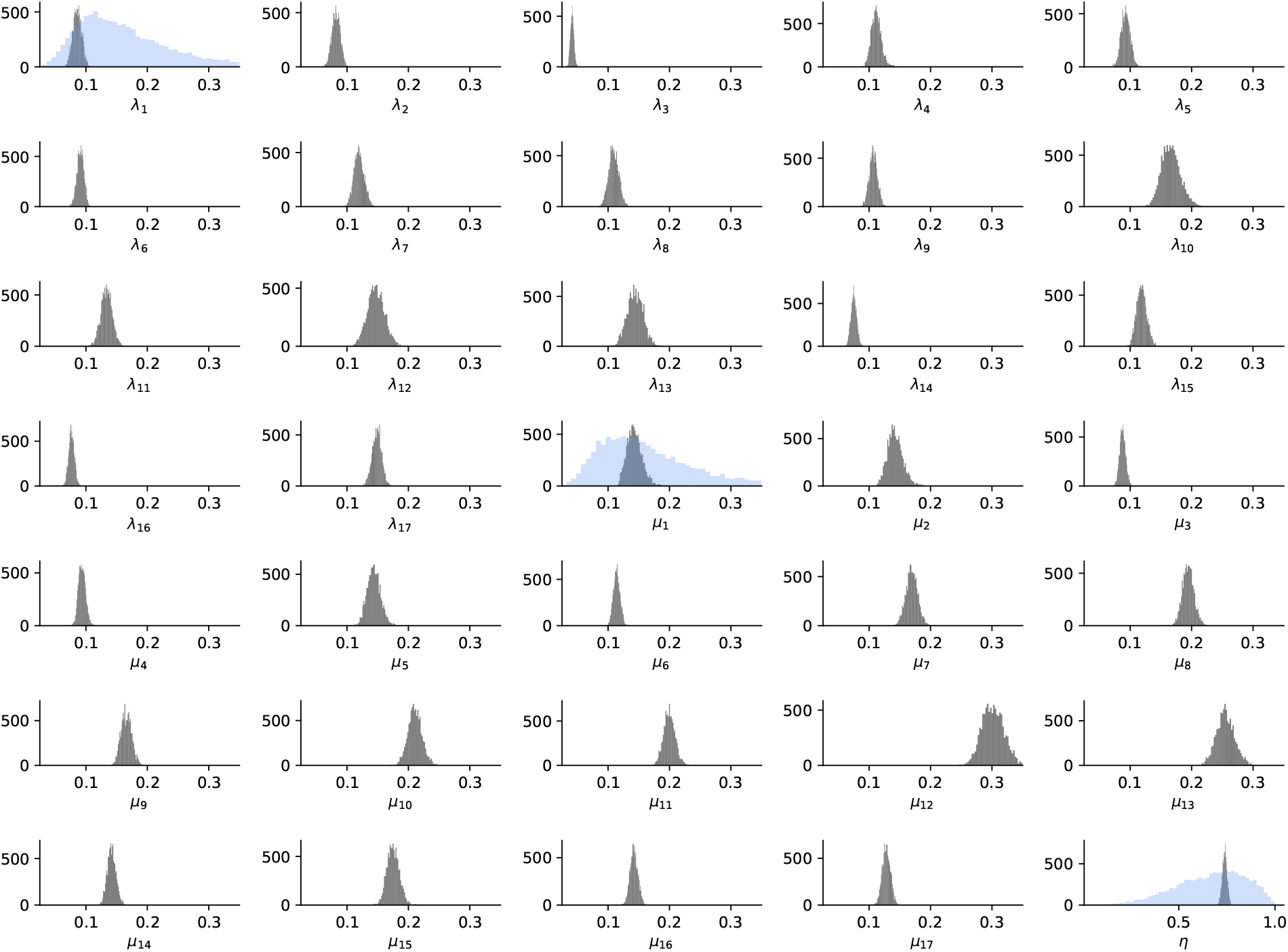
Posterior distributions for all duplication (*λ*) and loss *μ*) rates and the expected number of lineages at the root (1 /η) for the nine taxon analysis using the GBM prior with *v* = 0.1. The blue histograms denote the prior distributions on the rates at the root (*λ*_1_ and *μ*_1_) and the hyperprior on *η*.

**Figure S8:**
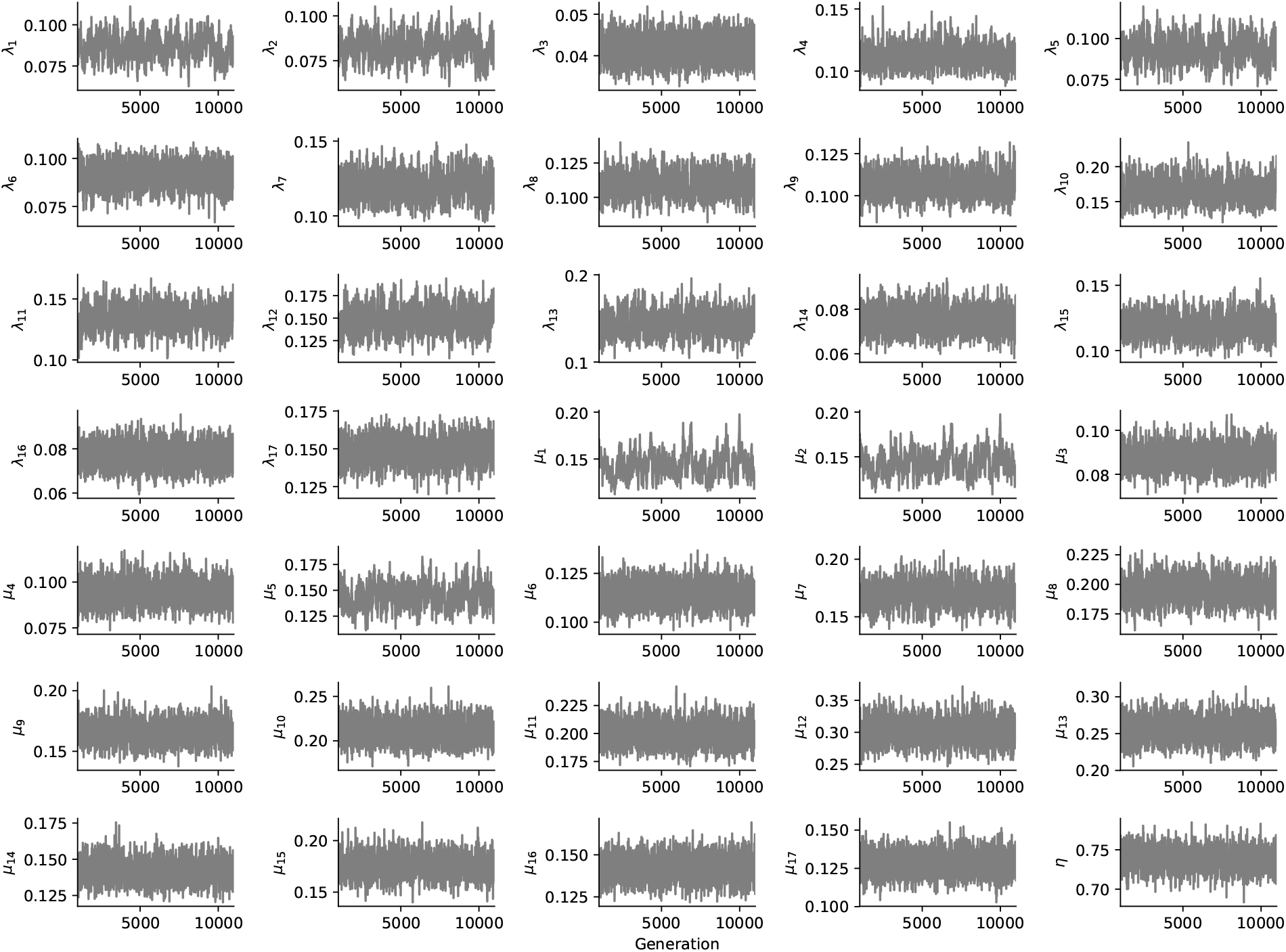
Trace plots for all duplication (*λ*) and loss (*μ*) rates and the expected number of lineages at the root (1/*η*) for the nine taxon analysis using the GBM prior with *v* = 0.1. 10,000 generations are shown, after discarding 1000 generations as a burn-in.

**Figure S9:**
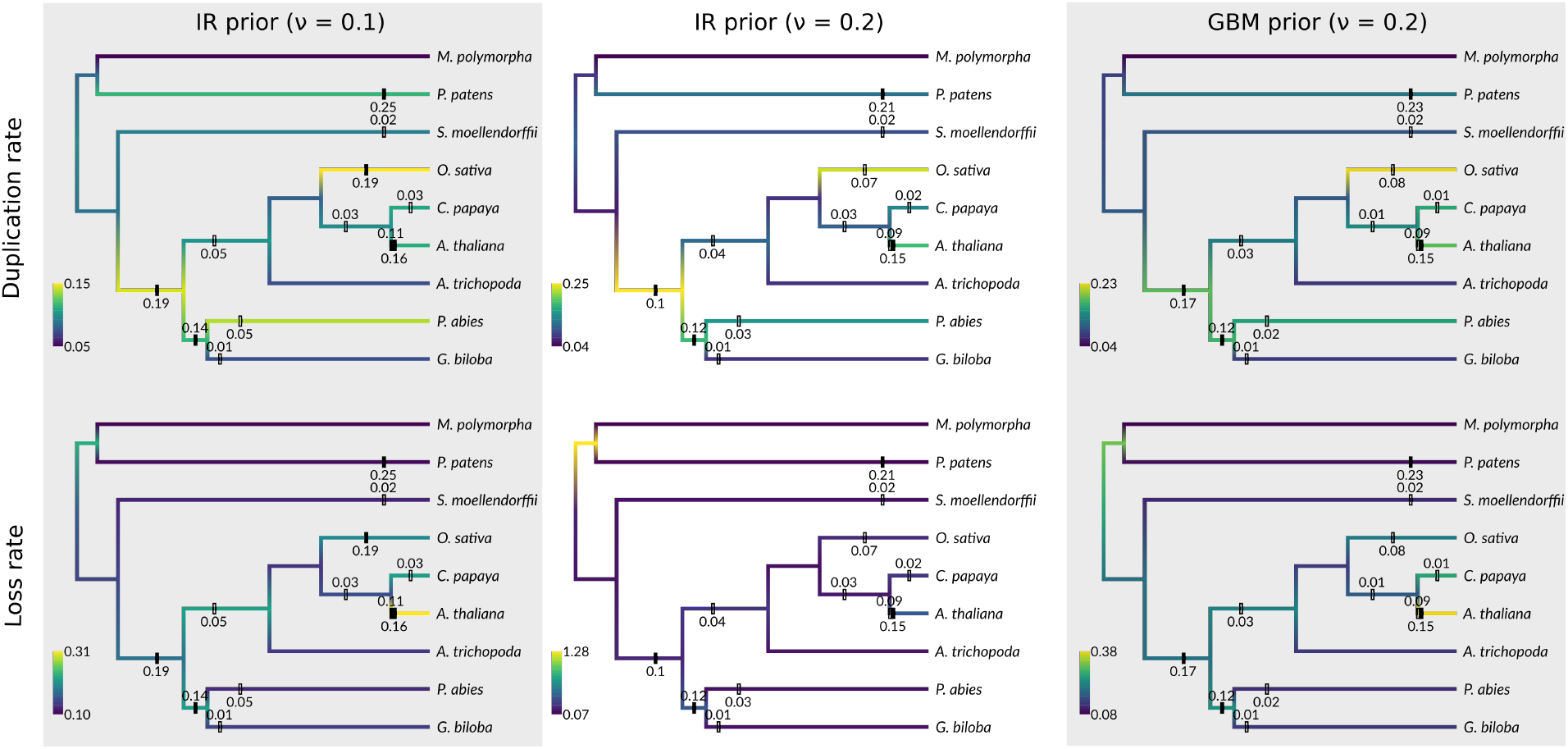
Duplication and loss rate estimates for the nine taxon data set under different priors. The colors indicate the mean of the marginal posterior for the relevant branch-wise rates.

**Figure S10:**
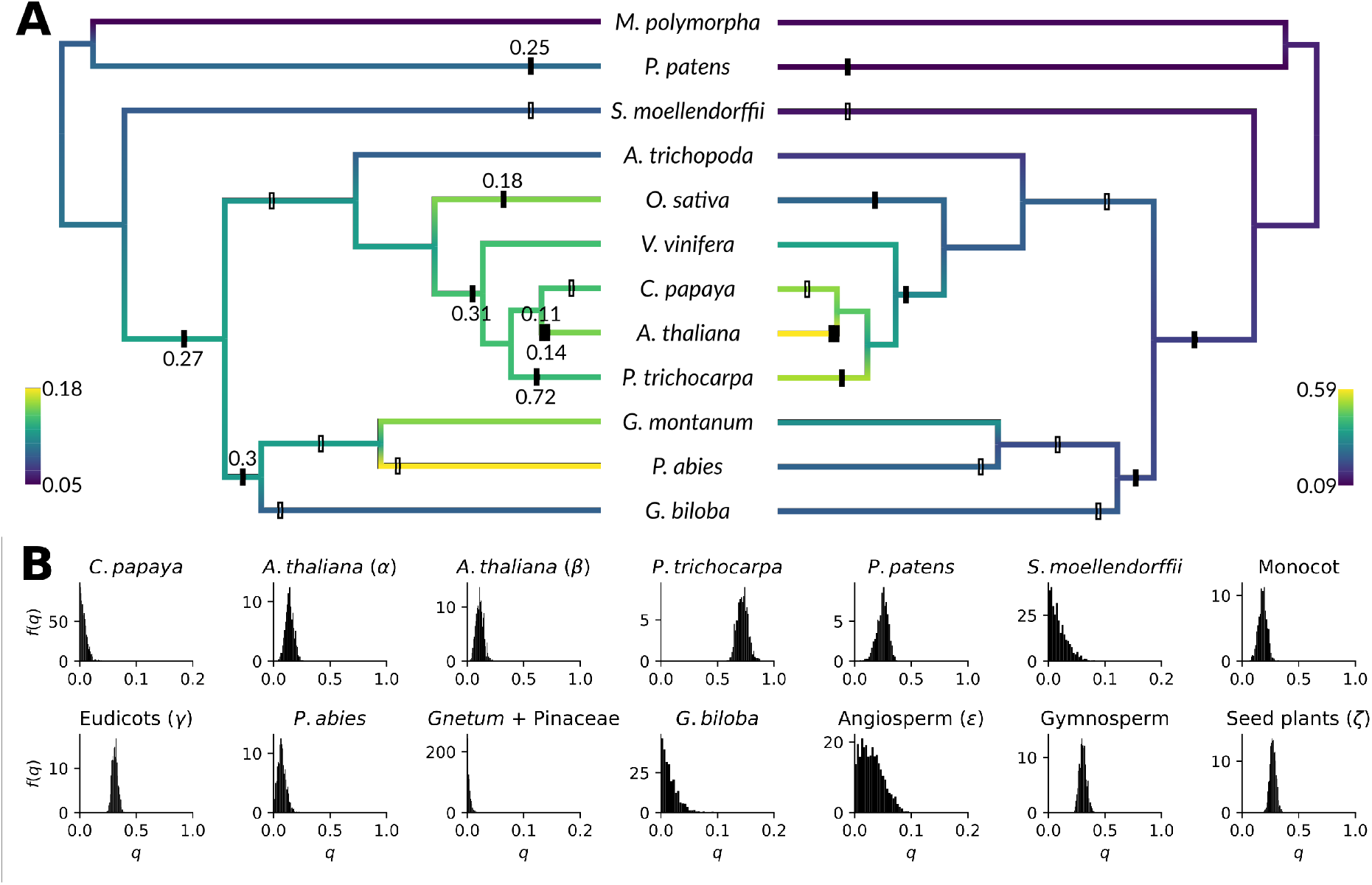
Duplication, loss and retention rate rate estimates for a 12 taxon data set, consisting of the union of the nine taxon, six taxon and five taxon data set, excluding *Pinus taeda* under a GBM prior on duplication and loss rates with *v* = 0.1. The colors indicate the mean of the marginal posterior for the relevant branch-wise rates while the bars indicate WGDs with there marginal posterior mean retention rate. Results are based on a sample of 5000 generations after a 1000 generation burn-in for 500 gene families.

**Figure S11:**
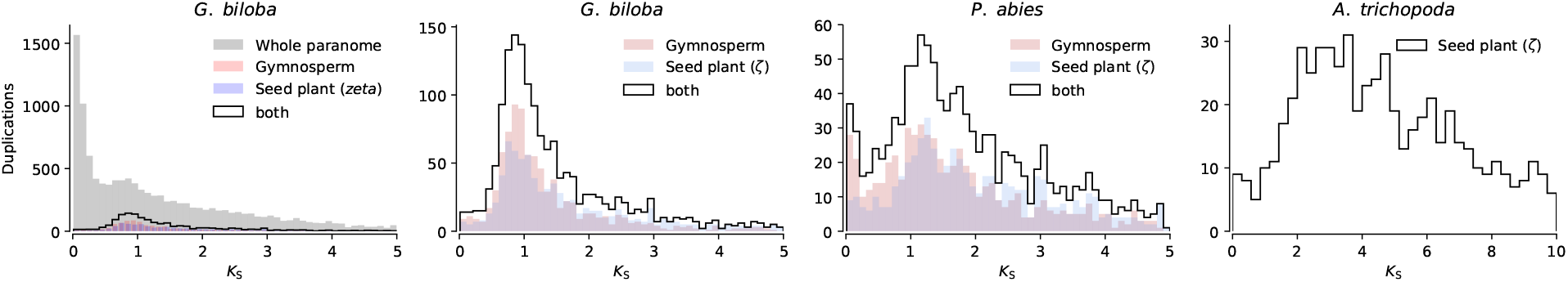
Node-averaged *K_S_* distributions for duplications reconciled to the hypothetical gymnosperm and seed plant WGD in *Ginkgo biloba, Picea abies* and *Amborella trichopoda* and the whole paranome *K_S_* distributions for *Ginkgo Biloba*.

**Figure S12:**
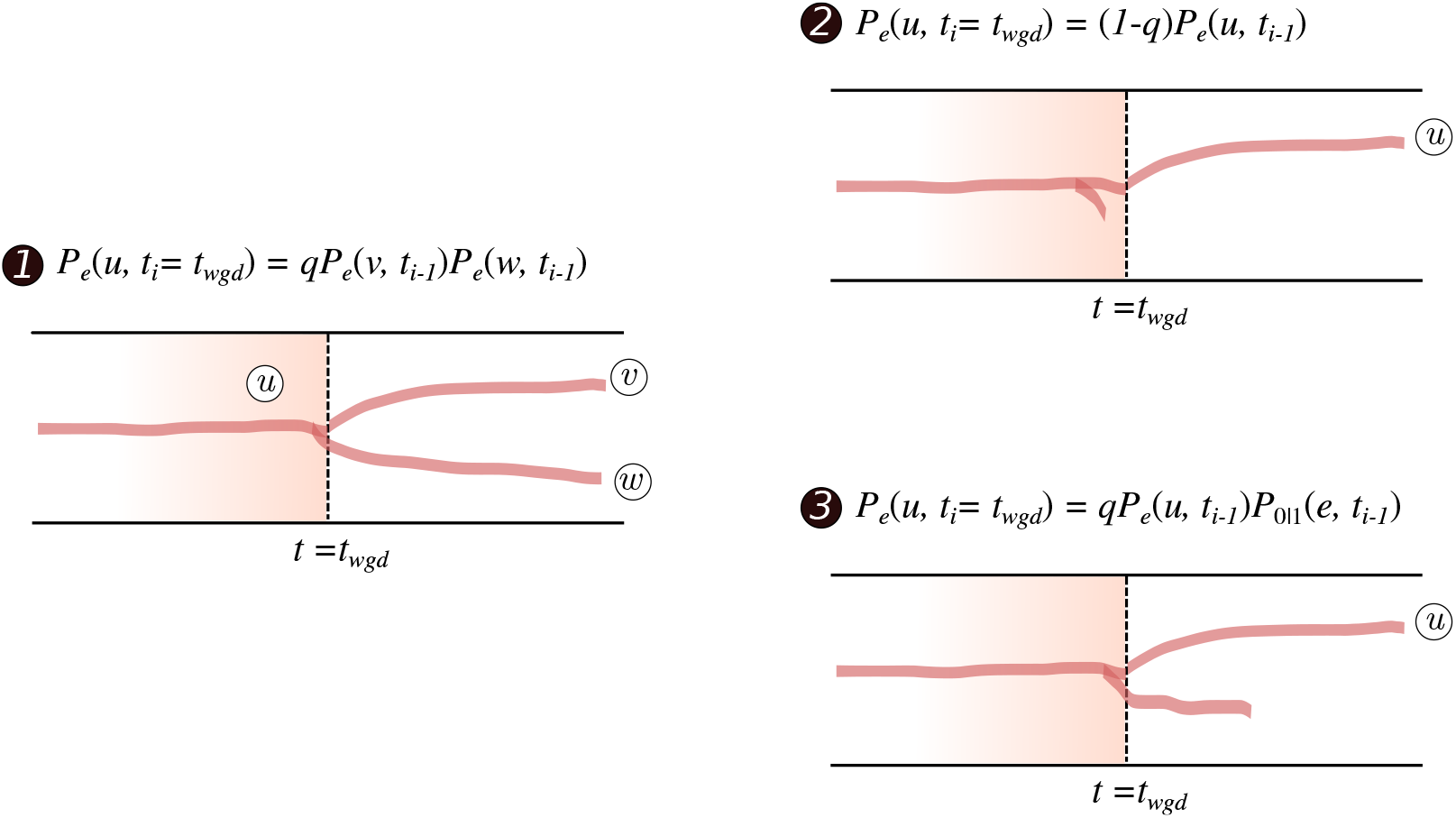
Illustration of the different scenario’s in the recursion at a WGD node. Each panel represents a species tree branch with a WGD node in the middle (dashed line). The time point of the WGD node coincides with the moment where rediploidization is complete. The color gradient illustrates the tetraploid phase where the WGD derived duplication has not yet been lost by rediploidization nor fixed at a new locus. Scenario (1) shows a retained WGD (with probability *q*), where a lineage leading to node *u* is retained after the WGD and leads to two daughter lineages *v* and *w*. Scenario (2) shows a non-retained WGD-derived duplication (with probability 1 − *q*), where during the diploidization process one of the duplicate copies was lost. Scenario (3) shows the case where one copy of an initially retained WGD-derived duplication is lost later on (at rate *μ*).

**Figure S13:**
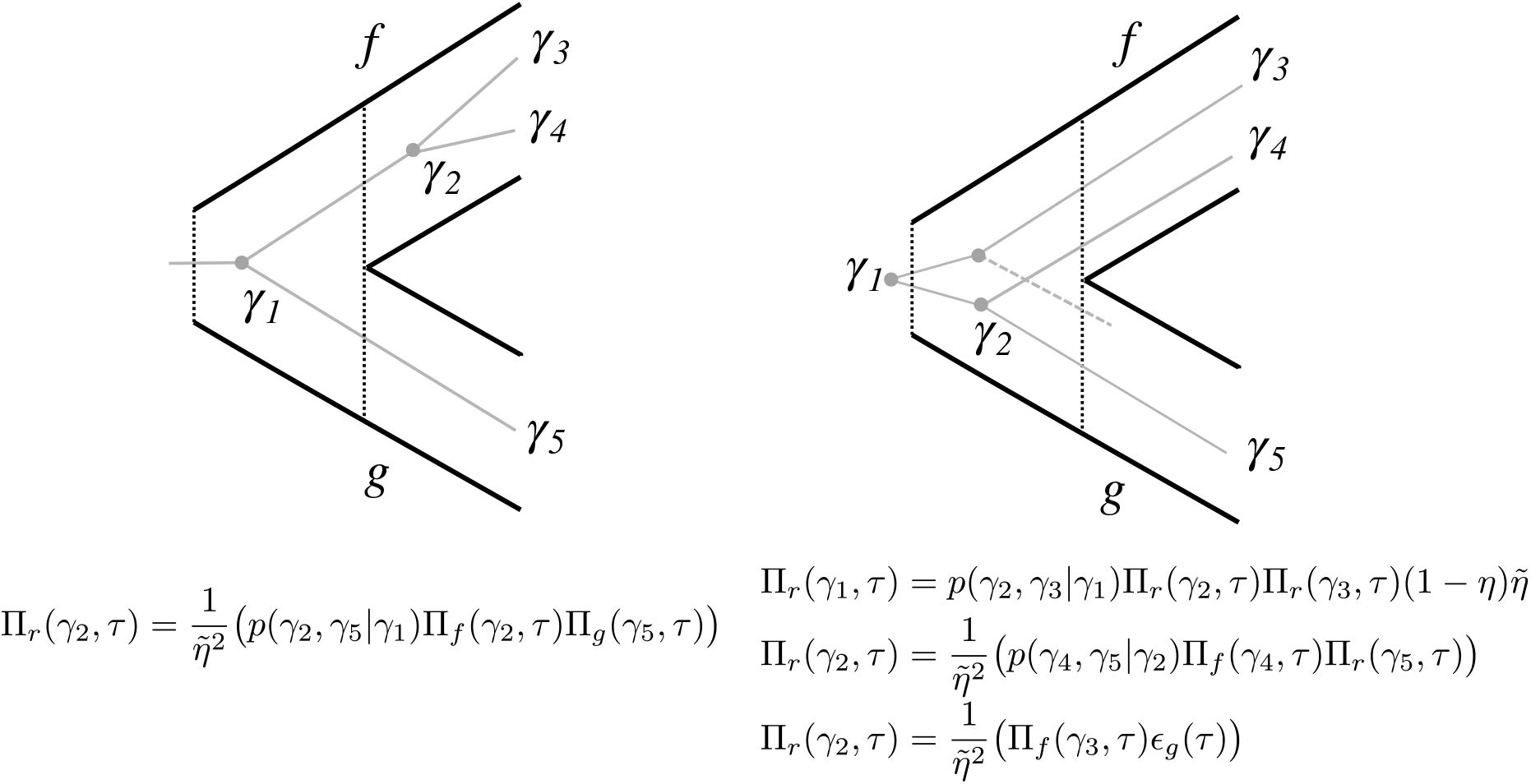
Illustration of the recursion at the root. In the example on the left there is one lineage entering the root, resulting in a prior probability of 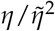. In the example on the right, two lineage enter the root of the species tree, resulting in a prior probability of 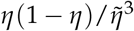. Note that Π_*τ*_(*γ*_1_) is the probability of observing *γ*_1_ at the root up to a factor *η*.

**Figure S14:**
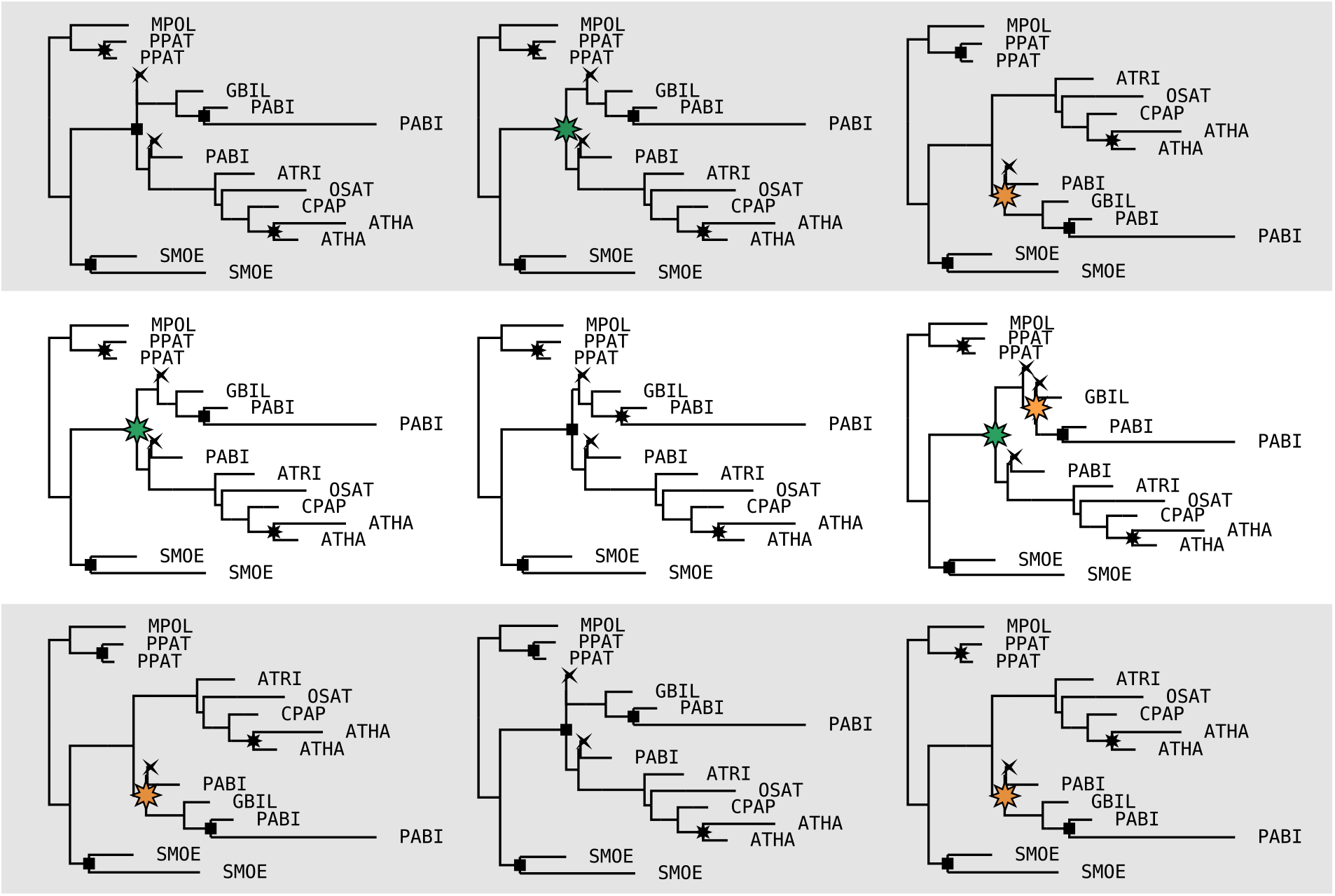
Nine reconciled trees sampled from the posterior distribution (using a GBM prior with *v* = 0.1) for the same gene family. The variation across individual trees illustrates how Whale deals with gene tree uncertainty through amalgamation. Duplication nodes are shown as squares, loss events as crosses and retained WGD events as stars. Orange stars highlight duplications reconciled to the hypothetical gymnosperm WGD whereas green stars highlight duplications reconciled tot the putative seed plant WGD. One can see that support for either one or both of these events is related to the gene tree topology.

**Figure S15:**
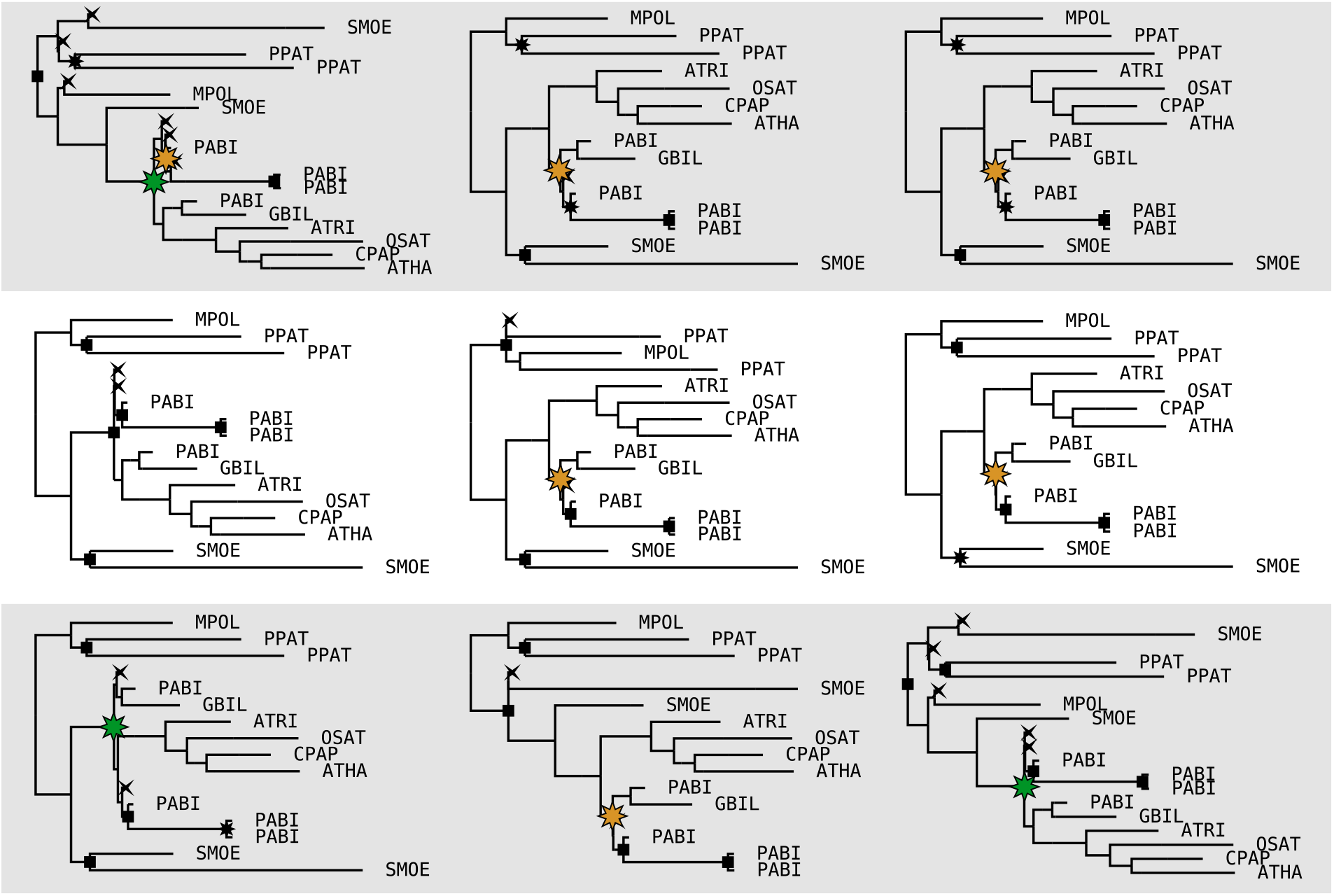
Nine reconciled trees sampled from the posterior distribution for the same gene family. Refer to Figure S14 for details.

**Figure S16:**
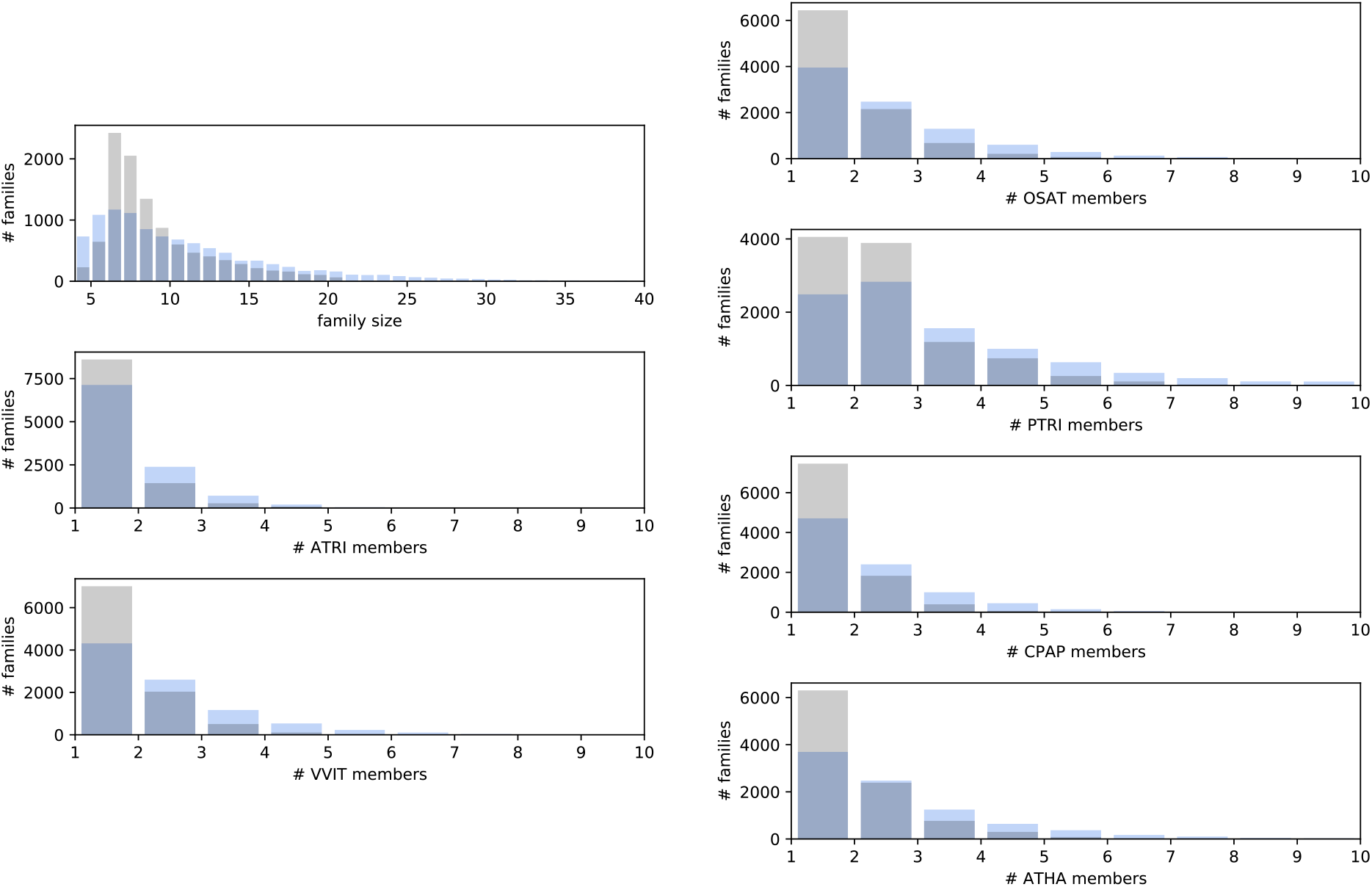
Posterior predictive checks for the six-taxon analysis. Posterior predictive checking was performed by simulating gene families from the joint posterior distribution inferred by MCMC, and comparing the distribution of gene family sizes and per-species gene counts with the observed (input) data.

**Figure S17:**
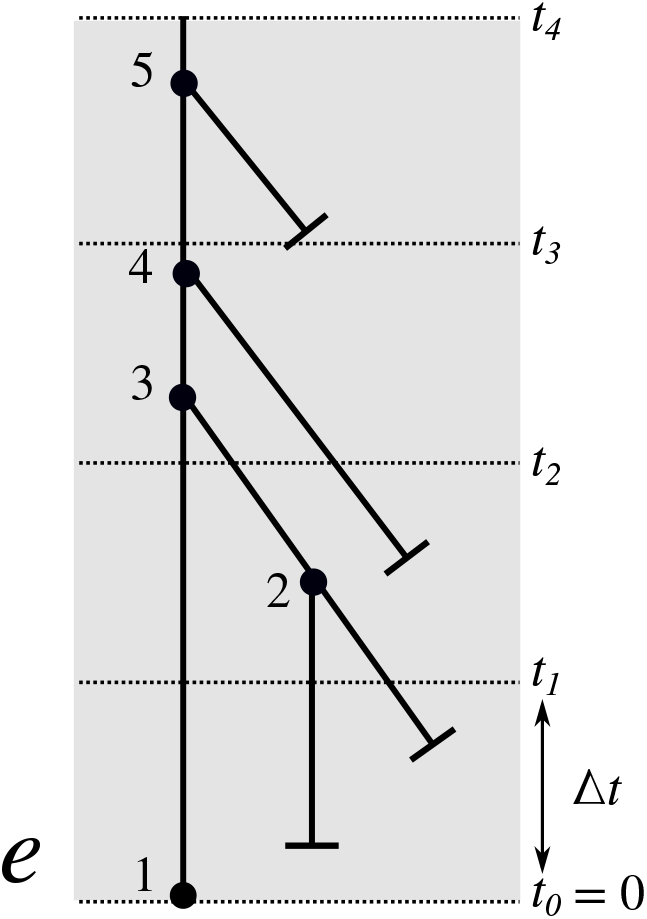
Ilustration of a branch *e* of the species tree discretized in multiple time slices of size Δ*t*. The extinction probability *ϵ_e_*(*t* + Δ*t*) at some time *t* + Δ*t* is simply the probability that the gene leaves no descendants observed at *t*0, which can be decomposed into the probability of extinction within the slice directly below *t* + Δ*t*, which is *α*(Δ*t*) and the probability that it does not go extinct, but does so in the part of the branch below the current time slice, which is 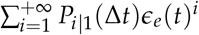. This last series can be further simplified to obtain equation (23). The propagation probability *ϕ*(*t, t* + Δ*t*) is the probability that a lineage at time *t* + Δ*t* gives rise to exactly one lineage at time *t* that has observed descendants. An example of such a scenario is the following: the lineage entering the time slice between *t*_2_ and *t*_3_ at *t*_3_ gives rise to three lineages at the end of this time slice through two (unobserved) duplication events at node 3 and 4. Of these three lineages eventually only one gives rise to observed descendants. The probability of such an event is 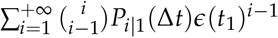, which simply considers all possible duplication-loss scenarios in a slice of length Δ*t* that give rise to *i* lineages at the end of the time slice (in this example *i* = 3) of which we pick *i* − 1 random lineages that go extinct further down in the tree. This infinite series can again be simplified to equation (24).

## References

Abbasi AA. 2010. Piecemeal or big bangs: Correlating the vertebrate evolution with proposed models of gene expansion events. Nature Reviews Genetics. 11(2):166.

Arvestad L, Lagergren J, Sennblad B. 2009. The gene evolution model and computing its associated probabilities. Journal of the ACM. 56(2):1–44.

Bailey NTJ. 1964. The Elements of Stochastic Processes with Applications to the Natural Sciences. New York-London: Wiley.

Blanc G, Wolfe KH. 2004. Widespread Paleopolyploidy in Model Plant Species Inferred from Age Distributions of Duplicate Genes. The Plant Cell. 16(7):1667–1678.

Brooks S, Gelman A, Jones G, Meng X-L. 2011. Handbook of markov chain monte carlo. CRC press.

Chen K, Durand D, Farach-Colton M. 2000. NOTUNG: A program for dating gene duplications and optimizing gene family trees. Journal of Computational Biology: A Journal of Computational Molecular CeΠ Biology. 7(3-4):429–447.

Clark JW, Donoghue PCJ. 2017. Constraining the timing of whole genome duplication in plant evolutionary history. Proceedings of the Royal Society B: Biological Sciences. 284(1858):20170912.

Csűirös M, Miklós I. 2009. Streamlining and large ancestral genomes in archaea inferred with a phylogenetic birth-and-death model. Molecular biology and evolution. 26(9):2087–2095.

Dickey JM. 1971. The weighted likelihood ratio, linear hypotheses on normal location parameters. The Annals of Mathematical Statistics.:204–223.

Emms DM, Kelly S. 2015. OrthoFinder: Solving fundamental biases in whole genome comparisons dramatically improves orthogroup inference accuracy. Genome biology. 16(1):157.

Guan R, Zhao Y, Zhang H, Fan G, Liu X, Zhou W, Shi C, Wang J, Liu W, Liang X, et al. 2016. Draft genome of the living fossil Ginkgo biloba. GigaScience. 5(1).

Hahn MW. 2007. Bias in phylogenetic tree reconciliation methods: Implications for vertebrate genome evolution. Genome biology. 8(7):R141.

Hahn MW, De Bie T, Stajich JE, Nguyen C, Cristianini N. 2005. Estimating the tempo and mode of gene family evolution from comparative genomic data. Genome Research. 15(8):1153–1160.

Höhna S, Drummond AJ. 2012. Guided Tree Topology Proposals for Bayesian Phylogenetic Inference. Systematic Biology. 61(1):1–11.

Jiao Y, Leebens-Mack J, Ayyampalayam S, Bowers JE, McKain MR, McNeal J, Rolf M, Ruzicka DR, Wafula E, Wickett NJ, et al. 2012. A genome triplication associated with early diversification of the core eudicots. Genome biology. 13(1):R3.

Jiao Y, Li J, Tang H, Paterson AH. 2014. Integrated syntenic and phylogenomic analyses reveal an ancient genome duplication in monocots. The Plant Cell.:tpc-114.

Jiao Y, Wickett NJ, Ayyampalayam S, Chanderbali AS, Landherr L, Ralph PE, Tomsho LP, Hu Y, Liang H, Soltis PS, et al. 2011. Ancestral polyploidy in seed plants and angiosperms. Nature. 473(7345):97–100.

Kishino H, Hasegawa M. 1990. Converting distance to time: Application to human evolution.

Larget B. 2013. The Estimation of Tree Posterior Probabilities Using Conditional Clade Probability Distributions. Systematic Biology. 62(4):501–511.

Li F-W, Brouwer P, Carretero-Paulet L, Cheng S, De Vries J, Delaux P-M, Eily A, Koppers N, Kuo L-Y, Li Z, et al. 2018. Fern genomes elucidate land plant evolution and cyanobacterial symbioses. Nature plants. 4(7):460.

Li Z, Baniaga AE, Sessa EB, Scascitelli M, Graham SW, Rieseberg LH, Barker MS. 2015. Early genome duplications in conifers and other seed plants. Science Advances. 1(10):e1501084.

Li Z, Defoort J, Tasdighian S, Maere S, Van de Peer Y, De Smet R. 2016. Gene Duplicability of Core Genes Is Highly Consistent across All Angiosperms. The Plant Cell. 28(2):326–344.

Li Z, Tiley GP, Galuska SR, Reardon CR, Kidder TI, Rundell RJ, Barker MS. 2018. Multiple large-scale gene and genome duplications during the evolution of hexapods. Proceedings of the National Academy of Sciences. 115(18):4713–4718.

Li Z, Tiley GP, Rundell RJ, Barker MS. 2019. Reply to nakatani and mclysaght: Analyzing deep duplication events. Proceedings of the National Academy of Sciences.:201819227.

Loytynoja A, Goldman N. 2008. Phylogeny-aware gap placement prevents errors in sequence alignment and evolutionary analysis. Science. 320(5883):1632–1635.

Lynch M, Conery J. 2000. The Evolutionary Fate and Consequences of Duplicate Genes. Science. 290(5494):1151–1155.

Maere S, De Bodt S, Raes J, Casneuf T, Van Montagu M, Kuiper M, Van de Peer Y. 2005. Modeling gene and genome duplications in eukaryotes. Proceedings of the National Academy of Sciences. 102(15):5454–5459.

McKain MR, Tang H, McNeal JR, Ayyampalayam S, Davis JI, dePamphilis CW, Givnish TJ, Pires JC, Stevenson DW, Leebens-Mack JH. 2016. A Phylogenomic Assessment of Ancient Polyploidy and Genome Evolution across the Poales. Genome Biology and Evolution. 8(4):1150–1164.

Morris JL, Puttick MN, Clark JW, Edwards D, Kenrick P, Pressel S, Wellman CH, Yang Z, Schneider H, Donoghue PC. 2018. The timescale of early land plant evolution. Proceedings of the National Academy of Sciences. 115(10):E2274–E2283.

Nakatani Y, McLysaght A. 2019. Macrosynteny analysis shows the absence of ancient whole-genome duplication in lepidopteran insects. Proceedings of the National Academy of Sciences.:201817937.

Nystedt B, Street NR, Wetterbom A, Zuccolo A, Lin Y-C, Scofield DG, Vezzi F, Delhomme N, Giacomello S, Alexeyenko A, et al. 2013. The Norway spruce genome sequence and conifer genome evolution. Nature. 497(7451):579–584.

Rabier C-E, Ta T, Ané C. 2014. Detecting and Locating Whole Genome Duplications on a Phylogeny: A Probabilistic Approach. Molecular Biology and Evolution. 31(3):750–762.

Rambaut A, Bromham L. 1998. Estimating divergence dates from molecular sequences. Molecular biology and evolution. 15(4):442–448.

Rannala B, Yang Z. 2007. Inferring Speciation Times under an Episodic Molecular Clock. Systematic Biology. 56(3):453–466.

Rasmussen MD, Kellis M. 2011. A Bayesian Approach for Fast and Accurate Gene Tree Reconstruction. Molecular Biology and Evolution. 28(1):273–290.

Roberts GO, Rosenthal JS. 2009. Examples of Adaptive MCMC. Journal of Computational and Graphical Statistics. 18(2):349–367.

Robertson FM, Gundappa MK, Grammes F, Hvidsten TR, Redmond AK, Lien S, Martin SA, Holland PW, Sandve SR, Macqueen DJ. 2017. Lineage-specific rediploidization is a mechanism to explain time-lags between genome duplication and evolutionary diversification. Genome biology. 18(1):111.

Ronquist F, Teslenko M, Van Der Mark P, Ayres DL, Darling A, Höhna S, Larget B, Liu L, Suchard MA, Huelsenbeck JP. 2012. MrBayes 3.2: Efficient bayesian phylogenetic inference and model choice across a large model space. Systematic biology. 61(3):539–542.

Roodt D, Lohaus R, Sterck L, Swanepoel RL, Van de Peer Y, Mizrachi E. 2017. Evidence for an ancient whole genome duplication in the cycad lineage. Chiang T-Y, editor. PLOS ONE. 12(9):e0184454.

Ruprecht C, Lohaus R, Vanneste K, Mutwil M, Nikoloski Z, Van de Peer Y, Persson S. 2017. Revisiting ancestral polyploidy in plants. Science Advances. 3(7):e1603195.

Salter LA. 2001. Complexity of the likelihood surface for a large dna dataset. Systematic biology. 50(6):970–978.

Smith JJ, Keinath MC. 2015. The sea lamprey meiotic map improves resolution of ancient vertebrate genome duplications. Genome research.

Szöllősi GJ, Boussau B, Abby SS, Tannier E, Daubin V. 2012. Phylogenetic modeling of lateral gene transfer reconstructs the pattern and relative timing of speciations. Proceedings of the National Academy of Sciences. 109(43):17513–17518.

Szöllősi GJ, Rosikiewicz W, Boussau B, Tannier E, Daubin V. 2013. Efficient exploration of the space of reconciled gene trees. Systematic biology. 62(6):901–912.

Szöllősi GJ, Tannier E, Daubin V, Boussau B. 2015. The Inference of Gene Trees with Species Trees. Systematic Biology. 64(1):e42–e62.

Szöllősi GJ, Tannier E, Lartillot N, Daubin V. 2013. Lateral gene transfer from the dead. Systematic biology. 62(3):386–397.

Tasdighian S, Van Bel M, Li Z, Van de Peer Y, Carretero-Paulet L, Maere S. 2017. Reciprocally retained genes in the angiosperm lineage show the hallmarks of dosage balance sensitivity. The Plant Cell. 29(11):2766–2785.

Thomas WG, Ather SH, Hahn MW. 2017. Gene-tree reconciliation with mul-trees to resolve polyploidy events. Systematic biology. 66(6):1007–1018.

Thorne JL, Kishino H, Painter IS. 1998. Estimating the rate of evolution of the rate of molecular evolution. Molecular Biology and Evolution. 15(12):1647–1657.

Tiley GP, Ané C, Burleigh JG. 2016. Evaluating and Characterizing Ancient Whole-Genome Duplications in Plants with Gene Count Data. Genome Biology and Evolution. 8(4):1023–1037.

Tiley GP, Barker MS, Burleigh JG. 2018. Assessing the performance of ks plots for detecting ancient whole genome duplications. Genome biology and evolution. 10(11):2882–2898.

Tuskan GA, DiFazio S, Jansson S, Bohlmann J, Grigoriev I, Hellsten U, Putnam N, Ralph S, Rombauts S, Salamov A, et al. 2006. The genome of black cottonwood, populus trichocarpa (torr. & Gray). Science. 313(5793):1596–1604.

Van Bel M, Diels T, Vancaester E, Kreft L, Botzki A, Van de Peer Y, Coppens F, Vandepoele K. 2017. PLAZA 4.0: An integrative resource for functional, evolutionary and comparative plant genomics. Nucleic acids research. 46(D1):D1190–D1196.

Van de Peer Y, Maere S, Meyer A. 2010. 2R or not 2R is not the question anymore. Nature Reviews Genetics. 11(2):166.

Van de Peer Y, Mizrachi E, Marchal K. 2017. The evolutionary significance of polyploidy. Nature Reviews Genetics. 18(7):411–424.

Vanneste K, Baele G, Maere S, Van de Peer Y. 2014. Analysis of 41 plant genomes supports a wave of successful genome duplications in association with the Cretaceous–Paleogene boundary. Genome Research. 24(8):1334–1347.

Vanneste K, Van de Peer Y, Maere S. 2013. Inference of Genome Duplications from Age Distributions Revisited. Molecular Biology and Evolution. 30(1):177–190.

Verdinelli I, Wasserman L. 1995. Computing bayes factors using a generalization of the savage-dickey density ratio. Journal of the American Statistical Association. 90(430):614–618.

Wegrzyn JL, Lee JM, Tearse BR, Neale DB. 2008. TreeGenes: A forest tree genome database. International journal of plant genomics. 2008.

Yang Y, Moore MJ, Brockington SF, Mikenas J, Olivieri J, Walker JF, Smith SA. 2018. Improved transcriptome sampling pinpoints 26 ancient and more recent polyploidy events in Caryophyllales, including two allopolyploidy events. New Phytologist. 217(2):855–870.

Yang Z. 1994. Maximum likelihood phylogenetic estimation from DNA sequences with variable rates over sites: Approximate methods. Journal of Molecular Evolution. 39(3):306–314.

Yang Z. 1998. Likelihood ratio tests for detecting positive selection and application to primate lysozyme evolution. Molecular Biology and Evolution. 15(5):568–573.

Yoder AD, Yang Z. 2000. Estimation of primate speciation dates using local molecular clocks. Molecular Biology and Evolution. 17(7):1081–1090.

Zmasek CM, Eddy SR. 2001. A simple algorithm to infer gene duplication and speciation events on a gene tree. Bioinformatics (Oxford, England). 17(9):821–828.

Zwaenepoel A, Li Z, Lohaus R, Van de Peer Y. 2018. Finding evidence for whole genome duplications: A reappraisal. Molecular Plant.

Zwaenepoel A, Van de Peer Y. 2019. Wgd: Simple command line tools for the analysis of ancient whole genome duplications. Bioinformatics.

